# Hippo signaling regulates cuticle pigmentation and dopamine metabolism in *Drosophila*

**DOI:** 10.1101/2025.11.13.688152

**Authors:** Shelley B Gibson, Samantha L Deal, Ye-Jin Park, Bo Sun, Yanyan Qi, Jung-Wan Mok, Hyung-Lok Chung, Hongjie Li, Shinya Yamamoto

**Affiliations:** Department of Molecular and Human Genetics, Baylor College of Medicine (BCM), Houston, TX 77030, USA; Jan and Dan Duncan Neurological Research Institute (NRI), Texas Children’s Hospital (TCH), Houston, TX 77030, USA; Department of Biology, College of Liberal Arts & Sciences, Mercer University, Macon, GA 31207, USA; Development, Disease Models & Therapeutics Graduate Program, BCM, Houston, TX 77030, USA; Huffington Center on Aging, BCM, Houston, TX 77030, USA; Department of Neurology, Houston Methodist Research Institute, Houston, TX 77030 USA; Department of Neuroscience, BCM, Houston, TX 77030 USA

**Author notes:** Corresponding Author: Shinya Yamamoto; Fax: N/A.

**Keywords:** *Drosophila melanogaster*, dopamine, pigmentation, Hippo signaling, *warts*, *yorkie*

## Abstract

Pigmentation plays multiple important roles in development, physiology and evolution. Melanization of the insect cuticle requires dopamine as a precursor of melanin and involves key enzymes in dopamine biosynthesis including Tyrosine hydroxylase (TH) and Dopa decarboxylase (Ddc). Some studies have hinted that disruption of the evolutionarily conserved Hippo signaling pathway, which has been primarily studied in the context of tissue growth, may lead to changes in cuticle pigmentation in the fruit fly *Drosophila melanogaster*. However, to our knowledge, there have not been any systematic investigations into their potential mechanistic links. In this study, we identified that all genes that comprise the canonical Hippo signaling pathway [*hippo* (*hpo*), *salvador* (*sav*), *mats*, *warts* (*wts*), *yorkie* (*yki*) and *scalloped* (*sd*)] are involved in cuticle pigmentation in the fly notum based on tissue-specific gene knock-down/out experiments and epistatic analysis. Despite the notable divergence of pigmentation mechanisms between invertebrates and vertebrates, these phenotypes can often be rescued by the human orthologs of corresponding fly genes. While we find that manipulation of Hippo signaling in dopaminergic neurons does not affect global dopamine levels in the fly brain, developmental inhibition of this pathway can increase dopamine levels in the fly head, indicating that the mechanism by which dopamine levels are regulated in the nervous system is distinct from that in epithelial cells. Through single nuclei RNA sequencing of the developing fly nota and subsequent functional studies of differentially expressed genes that are altered upon inhibition of Hippo signaling, we found many genes that contribute to cuticle pigmentation downstream of Hippo signaling. We conclude that regulation of cuticle pigmentation by canonical Hippo signaling acts through multiple downstream genes rather than directly through *TH* and *Ddc.* We also propose that the *Drosophila melanogaster* cuticle may serve as a useful platform to identify previously uncharacterized mediators of Hippo signaling as well as an *in vivo* experimental system to test the functionality of rare genetic variants found in human Hippo signaling orthologs associated with a variety of diseases.

## Introduction

The Hippo signaling pathway is an integral pathway for growth and development and is classically known for its role in regulating cell proliferation, organ growth and apoptosis [1,2]. There are more than 30 proteins involved in the Hippo signaling pathway in the fruit fly *Drosophila melanogaster* [3], including Hippo (Hpo), Salvador (Sav), Mats (Mts), Warts (Wts), Yorkie (Yki) and Scalloped (Sd), that play key roles in the canonical kinase signaling cascade that leads to transcriptional modulation [4]. When Hippo signaling is activated, Hpo interacts with Sav to phosphorylate the Wts/Mats complex which in turn phosphorylates Yki [5–9]. This phosphorylation event prevents Yki translocation into the nucleus and subsequent Yki-mediated co-transcriptional activation function, primarily through Sd (**Fig 1A**, [10–15]). This process is considered as the ‘canonical Hippo signaling pathway’, but there are several modes of regulation at different entry points of the pathway components upstream of Yki repression as well as direct regulation of Yki/Sd outside of Hippo signaling [3,16].

**Fig 1.**
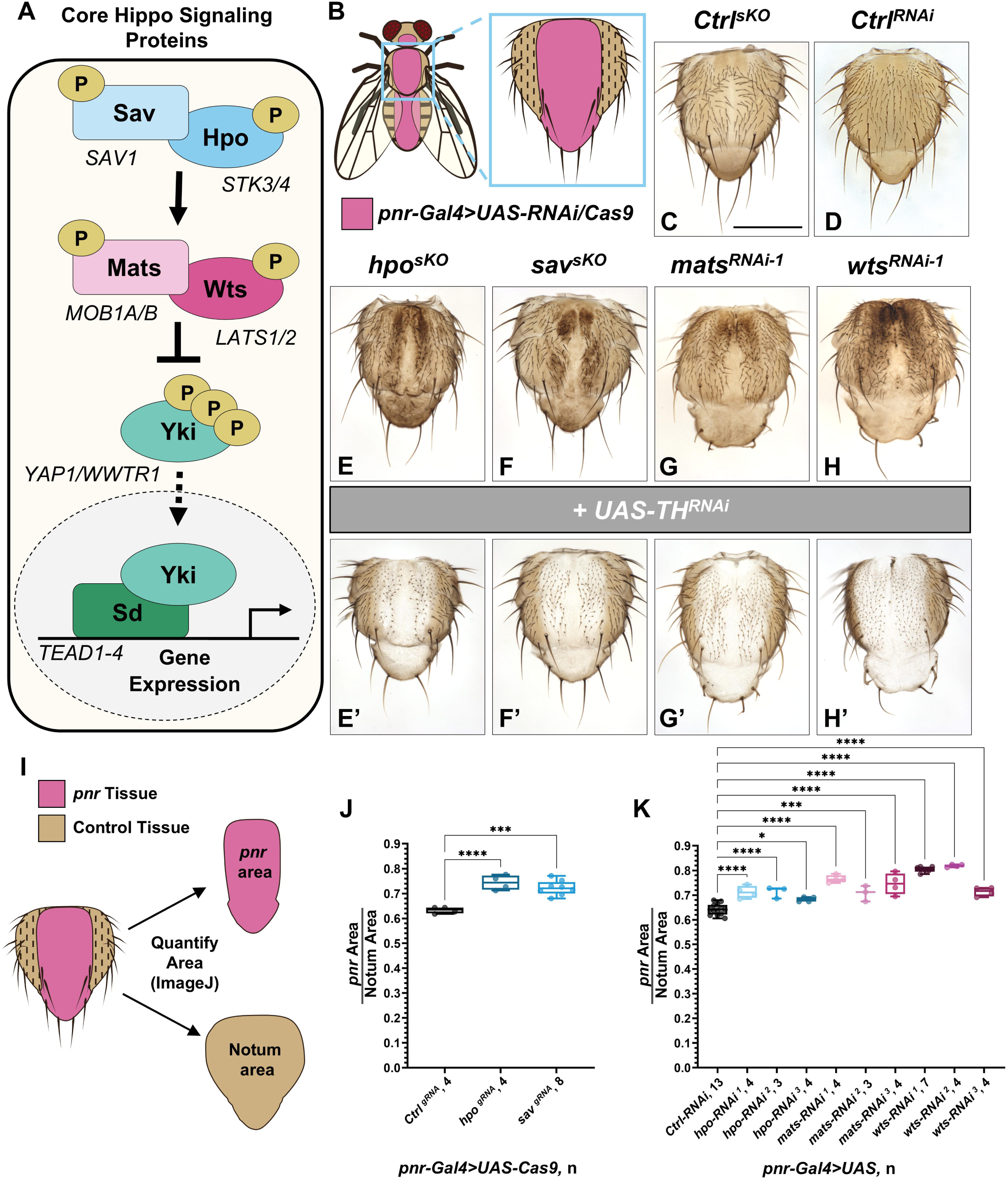
Inhibiting core Hippo signaling genes leads to cuticle overgrowth and increased pigmentation. **(A)** The core canonical Hippo signaling pathway in *Drosophila melanogaster* in bold with human orthologs listed underneath. **(B)** Domain pattern of *pannier*(*pnr*)*-Gal4* driving UAS elements in the adult fly [147]. **(C)** Dissected adult notum images of control (Ctrl) somatic CRISPR gRNA knockout (sKO) flies and **(D)** RNAi knockdown (KD) flies. **(E-F)** Increased pigmentation and overgrowth growth phenotypes with sKO of (**E**) *hpo* and **(F)** *sav* using gRNAs. **(G-H)** Similar phenotypes are seen with RNAi KD of **(G)** *mats* and **(H)** *wts*. **(E’-H’)** Suppression of pigmentation, but not growth with co-KD of *TH* using *UAS-TH-RNAi*. All experiments were performed at 29°C. Scale bar = 0.5mm. **(I)** Schematic of areas quantified to determine proportion of notum that is expressing *pnr-Gal4*, designated as *pnr* area. **(J)** Area quantification of sKO of *hpo* and *sav*. Statistical significance was assessed by one-way ANOVA pair-wise comparison to *Ctrl* (n=4-8, 29°C). **(K)** Area quantification of RNAi KD of *hpo*, *mats* and *wts*. Statistical significance was determined by one-way ANOVA pair-wise compared to *Ctrl* (n=3-13, 29°C).

Canonical and some non-canonical components of Hippo signaling as well as their biological functions are highly conserved across species including mammals [3,4,16]. Hippo signaling regulation is important for human health due to its roles in development, tissue homeostasis and regeneration [17,18]. Disruption of this pathway leads to severe problems in several organ systems [19–28] as well as the immune system [29–31]. Ultimately, the most common and characterized human disease observed with Hippo signaling problems are several different types of cancer [32–35]. This uncontrolled growth phenotype is also observed in flies as well, for example when mutant clones of the upstream genes are induced or *yki* is overexpressed [7,36–39]. Hippo signaling has also been implicated in disorders that affect the nervous system [40–43], highlighting the need to study the molecular function of these genes in the nervous system and other relevant contexts.

Through a large scale RNA interference (RNAi) screen that was used to find novel regulators of dopamine levels, we identified that knockdown (KD) of three genes in the Hippo signaling pathway (*hpo*, *mats* and *wts*) causes increased pigmentation of the *Drosophila* cuticle [44]. Although it has been hypothesized that in humans Hippo signaling regulates the development and maintenance of melanocytes [45,46], the cells responsible for skin pigmentation in humans, to our knowledge there has been no direct mechanistic studies that investigated the biological links between Hippo signaling and pigmentation in the fly. The mechanisms by which pigmentation is regulated in fly and human are not considered to be conserved as insect cuticle pigmentation is regulated primarily by ectoderm-derived epithelial cells that arise and develop differently than mammalian neural crest-derived melanocytes [47–49]. In addition, there are fundamental differences in the biochemical pathways that produce pigments between human and insects [50–52]. Importantly in *Drosophila*, melanin is made using the same biosynthetic enzymes, TH and Ddc, that are used to generate dopamine in the brain (**Fig S1A**). While the mechanism of skin/cuticle pigmentation is different between vertebrates and invertebrates, the study of cuticle pigmentation has implications into diverse aspects of basic biology such as evolution [53] and may provide insights into dopamine/neuromelanin regulation in the mammalian brain [54,55]. In this study we aimed to determine how Hippo signaling regulates cuticle pigmentation during development and further assessed whether this pathway may impact dopamine levels in the brain in flies.

Here, we show that canonical Hippo signaling regulates cuticle pigmentation through Yki and Sd. Although key Hippo signaling genes are expressed in dopaminergic neurons, we found that inhibiting Hippo signaling increases dopamine levels in the fly cuticle but not in the brain, suggesting that dopamine metabolism in these two organ systems is regulated differently. To identify downstream target genes that contribute to pigment regulation when this pathway is altered, we performed single-nucleus RNA-sequencing (snRNA-seq) on *wts* knockdown (KD) animals at a critical time point for pigmentation. Through this approach, we identified that JNK signaling is a contributor to the pigmentation phenotype downstream of Hippo signaling. We also identified additional genes contributing to this phenotype and some possible genes involved in a negative feedback loop to fine-tune the phenotypic outcome. Based on these data, we argue that *Drosophila melanogaster* cuticle pigmentation is an underappreciated readout of Hippo signaling activity that provides new insights into its biological roles and further propose that this system can also be used to study the functional consequences of genetic variants that are found in patients.

## Results

### Inhibiting core Hippo signaling kinases leads to cuticle overgrowth and increased pigmentation

The canonical Hippo signaling pathway consists of genes that encode two adaptor proteins (*sav* and *mats*) and two kinases (*hpo* and *wts*) acting on the effector Yki through repressing phosphorylation marks (**Fig 1A** [5–8,14]). Previously, Mummery-Widmer et al., reported that knockdown of the upstream genes in the dorsomedial *pannier* (*pnr*) expressing tissue of the fly notum leads to overgrowth as well as increased pigmentation (**Fig 1B**, [56]). Furthermore, using independent RNAi lines, we validated these phenotypes for *hpo*, *mats* and *wts* (**Fig 1D**, **1G-1I**, **1K**) [44]. However, we were not able to obtain independent RNAi lines that exhibit this phenotype for *sav*.

To overcome this issue, we assessed whether we can use CRISPR-mediated somatic knockout (sKO) [57] to induce similar phenotypes based on the Gal4/UAS system [58] (**Fig 1B**). Using gRNAs targeted against *hpo* or *sav,* we observed that loss of these genes indeed cause increased pigmentation and overgrowth phenotypes (**Fig 1C**, **1E-1J**). Due to the clonal nature of sKO experiments, the pigmentation pattern is mosaic in the *pnr*-*Gal4* expressing region.

To demonstrate this darkening of notum observed upon sKO or KD of these genes are in fact a pigmentation increase and not a secondary phenotype associated with necrosis, we brought in an additional *UAS-TH-RNAi* to knockdown the rate limiting enzyme in the melanin synthesis (**Fig S1A**). This manipulation completely suppresses the darkened cuticle phenotypes, similar to *TH-RNAi* KD on its own (**Fig S1B-1C**), seen with the inhibition of all four upstream Hippo signaling genes, but does not suppress the overgrowth phenotypes (**Fig 1E’-1H’**). In conclusion, all upstream genes of the Hippo signaling pathway inhibit pigmentation in the fly nota.

### Hippo signaling regulation of cuticle overgrowth and pigmentation is mediated by *yorkie* and *scalloped*

Since canonical Hippo signaling acts through Yki, we next assessed whether this gene is also involved in cuticle pigmentation. We first overexpressed wild-type *UAS-yki* using *pnr-Gal4* and observed no change in the cuticle (**Fig 2A-2B**, **S2A**). However, when we simultaneously overexpressed the same construct while knocking down *wts* or *hpo*, we observed a worsening of both pigmentation and overexpression phenotypes (**Fig 2C-2D’**, **S2A**). Based on these results, we conclude that overexpression of Yki is not sufficient to induce any observable defects since its activity is tightly regulated by the upstream kinase complex in this specific context. Next, we used a transgenic line that allows us to overexpress a constitutively active mutant Yki (*UAS-Yki.S168A*) [14]. Since overexpression of this construct was lethal to the fly with *pnr-Gal4* even at 18°C, we performed this in a conditional manner by introducing a temperature sensitive *tub-Gal80[ts]* transgene to bypass lethality [59]. When we induced the expression of Yki.S168A in the third larval instar stage, we observed that this hyperactive Yki led to similar pigmentation and overgrowth phenotypes seen when inhibiting upstream Hippo signaling genes (**Fig 2E-2F**, **S2B**). Also similar to the previous experiments (**Fig 1E’-1H**’), inhibiting melanin synthesis by co-KD of *TH* suppressed the pigmentation phenotype induced by Yki.S168A expression (**Fig 2F**’). Based on these data, we conclude that gain-of-function (GOF) of Yki is sufficient to induce increased pigmentation.

**Fig 2.**
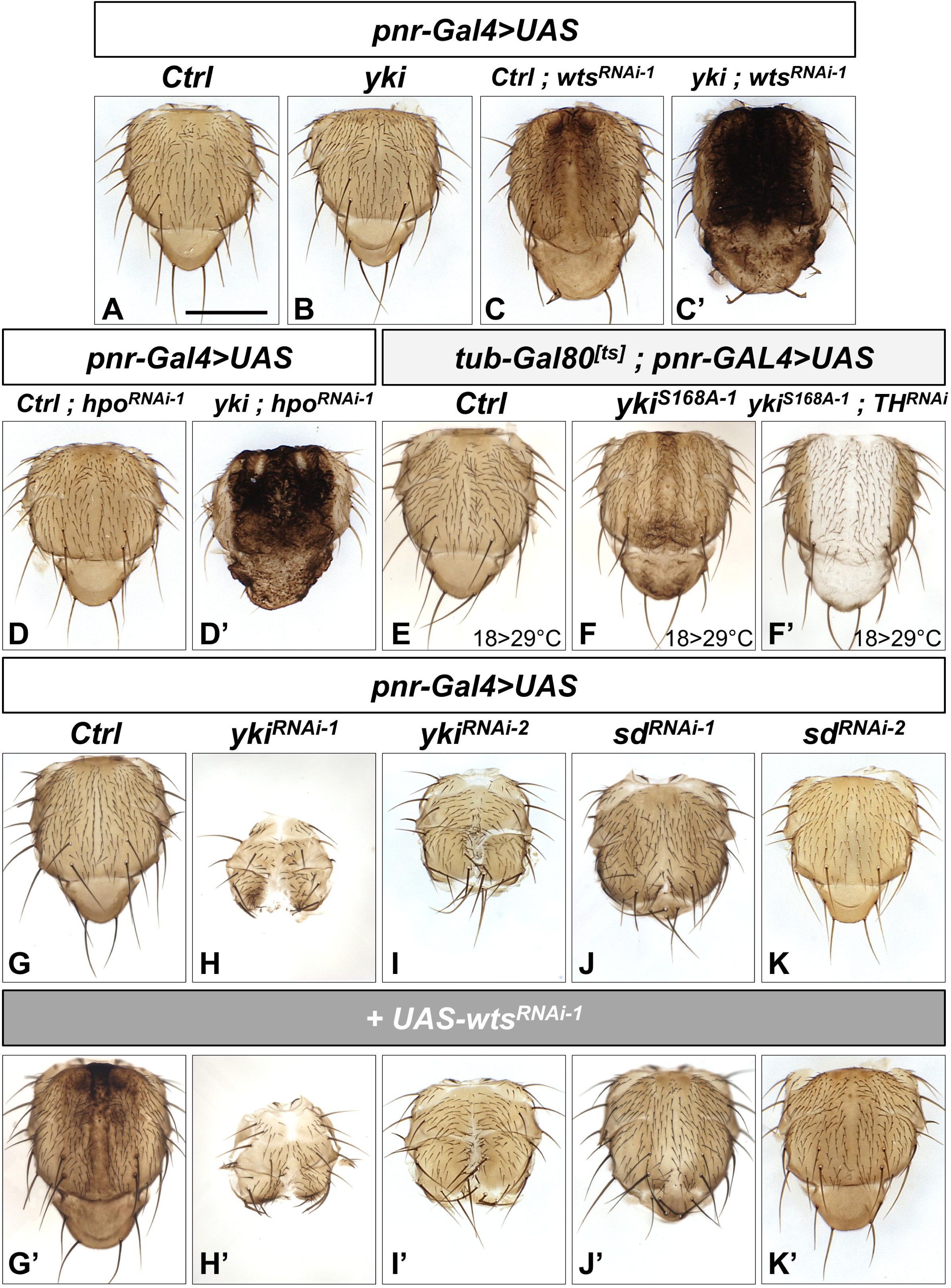
Hippo signaling regulates cuticle pigmentation through *yorkie* and *scalloped*. **(A)** Compared to *pnr>Ctrl*, **(B)** *yki* overexpression (OE) on its own does not show a phenotype. **(C)** While *wts* KD with a *Ctrl* OE construct gives an overgrowth and increased pigmentation phenotype, **(C’)** the *wts* KD phenotype is exacerbated with the addition of *yki* OE. **(D’)** *pnr*-Gal4 expression of *hpo*-RNAi has minimal phenotypes at 25°C, **(D’)** while adding *yki* OE leads to severe growth and pigmentation phenotypes. **(E)** Using temperature sensitive *tubGAL80[ts]* (18°C to 29°C switch in the wandering third instar larval stage), flies with expression of *pnr-Gal4>UAS-RNAi/OE,* control compared to **(F)** overactive Yki.S168A overexpression shows an overgrowth and increase in pigmentation. This increased pigmentation phenotype is suppressed by **(F’)** *TH-RNAi* KD. **(G)** Compared to a *pnr>Control(Ctrl)-RNAi,* **(H/I)** *yki* KD and **(J/K)** *sd* KD show strong undergrowth/dorsal closure phenotypes. **(G’)** *UAS-Ctrl-RNAi* with *wts* KD for epistatic analysis. KD of **(H’/I’)** *yki* and **(J’/K’)** *sd* are able to suppress and are epistatic to *wts* KD for both cuticle phenotypes. Grown at 25°C unless otherwise indicated. Scale bar = 0.5mm. Size quantification can be found in **Fig S2**.

To assess whether *yki* is necessary for cuticle pigmentation, we performed a KD experiment to determine whether loss-of-function (LOF) of this gene leads to decreased pigmentation. We observed a severe undergrowth phenotype upon *yki* KD in regions where *pnr-Gal4* is expressed (**Fig 2G-2I**). While this finding confirms that *yki* is necessary for organ size control in the nota, it was difficult to assess whether the LOF of this gene leads to reduced pigmentation due to the lack of tissues. We also did not quantify the undergrowth phenotype due to the severity of this phenotype. However, we were able to confirm that *yki* acts downstream of *wts* in this context based on epistasis experiments performed by co-KD of both genes, leading to the same severe phenotype seen with *yki* KD on its own (**Fig 2G’-I’**). Based on these data, we conclude that canonical Hippo signaling mediated by the Hpo/Sav-Mats/Wts-Yki axis regulates cuticle pigmentation in addition to tissue growth.

Yki is a transcriptional coactivator that requires transcription factors to regulate gene expression. In *Drosophila*, *sd* encodes the major transcription factor shown to mediate the function of Yki [10,11,13,60,61]. To determine whether Sd works with Yki to regulate these cuticle phenotypes, we performed additional KD and co-KD epistasis experiments (**Fig 2J-2K’**). When we knocked down *sd* with one of the two RNAi lines we obtained (line 1), we observed tissue undergrowth phenotypes that are similar to but milder than *yki* KD (**Fig 2J, S2C**). The second *sd* RNAi line we obtained produces no change in pigmentation and a slight undergrowth phenotype compared to control (**Fig 2K**, **S2C**). However, we found that *sd* KD mediated by both RNAi lines can strongly suppress both the overgrowth and pigmentation defects mediated by *wts* KD (**Fig 2J’-2K’, S2C**). This suggests that *sd* KD is epistatic to *wts* KD regarding both phenotypes. In conclusion, these results suggest that canonical Hippo signaling regulates cuticle pigmentation through *yki* and *sd*.

### Human orthologs of Hippo signaling genes can rescue cuticle phenotypes

While cuticle pigmentation is not a phenomenon that is conserved between flies and humans, all fly genes documented above have human orthologs. Therefore, we tested whether human counterparts of fly Hippo signaling genes can rescue the pigmentation defects in addition to the growth phenotype observed upon knocking down these fly genes. Typically, there is a one-to-two ratio of fly-to-human ortholog in the main components for the Hippo signaling pathways with the exception of the one-to-one ratio for *sav/SAV1* and one-to-four ratio for *Sd/TEAD1-4* (**Fig 1A**). In this study, we investigated the functional conservation of five human genes. KD of *mats* using *pnr-Gal4* leads to overgrowth and pigmentation phenotypes that we can rescue with human *MOB1A* or *MOB1B* expression **(Fig 3A-3C’, S3A)**. Similarly, we were able to rescue both growth and pigmentation phenotypes of *wts* KD with expression of human *LATS1* or *LATS2* (**Fig 3D-3F’, S3B**). We were also able to rescue the *yki* KD undergrowth phenotype with human *YAP1* (one of *yki*’s human orthologs, **Fig 3G-H’**). Note that the *yki* KD nota rescued by human *YAP1* has increased number of bristles (**Fig 3H’**), a phenotype we also observe when this gene is overexpressed in a wild-type thorax (**Fig 3H**). In conclusion, while cuticle pigmentation is not an evolutionarily conserved biological process, human orthologs of Hippo signaling pathway genes still retain the ability to regulate this process *in vivo*.

**Fig 3.**
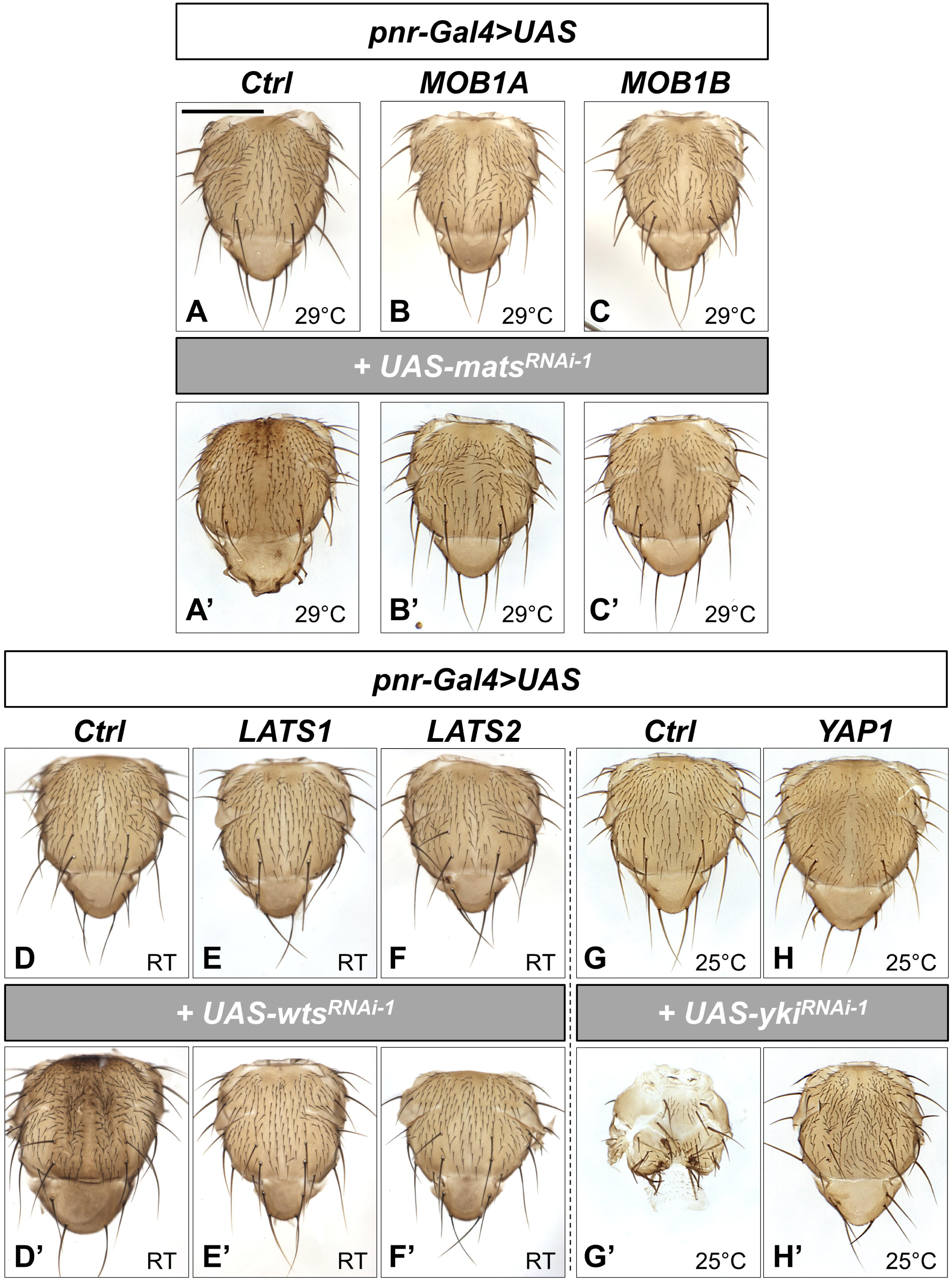
Multiple Hippo signaling human orthologs rescue cuticle growth and pigmentation phenotypes. OE of human cDNAs in the *pnr*-*Gal4* domain shows minimal phenotypes compared to **(A/D/G)** controls for **(B)** *MOB1A*, **(C)** *MOB1B*, **(E)** *LATS1*, and **(F)** *LATS2* at temperatures indicated. **(H)** *YAP1* OE shows an increase in bristle number at 25°C. **(A’)** *mats*-RNAi KD in a Ctrl OE background has a slightly darkened and overgrowth phenotype that are rescue by both of its human orthologs, **(B’)** *MOB1A* and **(C’)** *MOB1B*. **(D’, E’, F’)** Both *LATS1* and *LATS2* rescue the cuticle phenotypes associated with *wts* RNAi KD. **(G’)** KD of *yki* in the control OE background leads to similar undergrowth phenotypes seen previously, **(H’)** and this defect is rescued by expression of human *YAP1*. Growth temperatures are indicated in the figure image itself. Scale bar = 0.5mm. Size quantification can be found in **Fig S3**.

### Hippo signaling regulates global dopamine levels in the fly head

Cuticle pigmentation is primarily regulated by genes that are considered to affect dopamine synthesis and metabolism, some of which are evolutionarily conserved such as *TH* (encoded by the *pale* gene in *Drosophila*) and *Ddc* (**Fig S1A-C,** [50]). To determine if the increased pigmentation phenotype in the cuticle is accompanied by increase in dopamine levels, we performed High Performance Liquid Chromatography (HPLC) to measure dopamine levels in flies. As a positive control, we overexpressed TH, the rate-liming enzyme in dopamine synthesis, specifically in dopamine producing cells using *TH-Gal4* (also known as *ple-Gal4*) and confirmed that this manipulation leads to robust increase in dopamine levels extracted from fly heads (**Fig 4A**). When we inhibited Hippo signaling specifically in dopamine producing cells using *wts* RNAi driven by *TH-Gal4*, we also observed a significant increase in dopamine (**Fig 4A**), indicating that Hippo signaling regulates dopamine levels *in vivo*.

**Fig 4.**
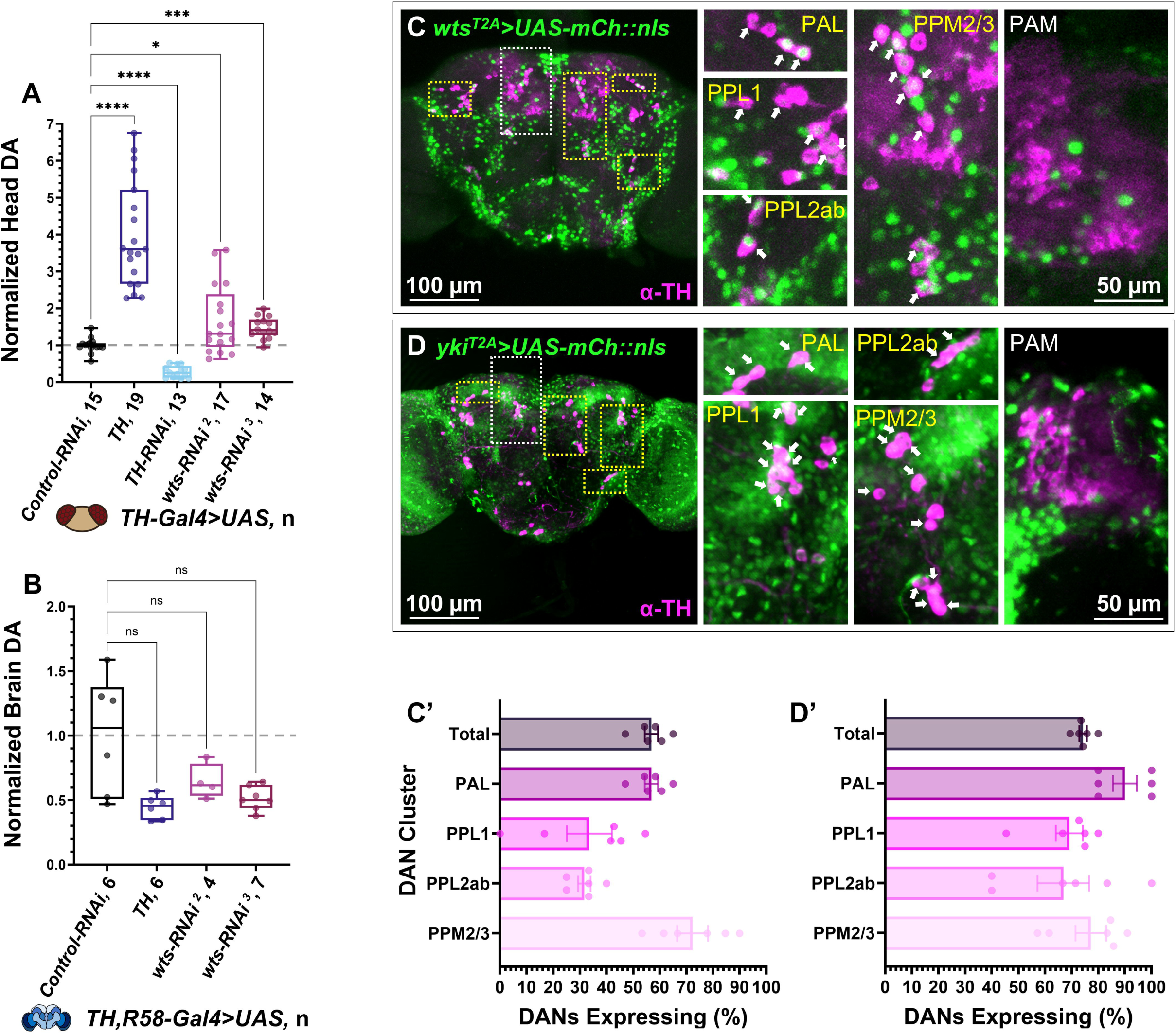
Hippo signaling regulates dopamine levels in the adult fly head but not in the brain. **(A)** High Performance Liquid Chromatography (HPLC) on adult fly heads expressing UAS elements via *TH*-*Gal4* show increase in dopamine (DA) levels with *wts*-RNAi KD compared to control (n=13-19, 5 heads/n, 29°C, 3-7 DAE). Statistical significance was determined by Brown-Forsythe and Welch’s ANOVA pair-wise analysis compared to control. **(B)** HPLC performed on brains with *wts*-RNAi KD show no significant difference from control (n=4-7, 10 brain/n, 29°C, 3-7 DAE). Statistical significance determined by Kruskal-Wallis pair-wise compared to control. **(C)** Full brain images of adult flies with *wts^T2A^* or **(D)** *yki^T2A^* driving expression of *UAS-mCherry::nls*. *T2A-Gal4>UAS-mCherry::nls* signal is shown in green while antibody TH staining is in magenta. Scale= 100µm. To the right of each are zoomed images of specific dopaminergic clusters, PAL, PPL1,PPL2ab and PPM2/3, outlined in yellow, while PAM on anterior side, outlined in white. Scale bar = 50µm. Filled white arrows show where TH+ expression overlaps in the same cell with T2A>mCh::nls (n= 3). Quantification of **(C’)** *wts^T2A^* and **(D’)** *yki^T2A^ >mCh::nls* expression in TH+ neurons, n= 6 hemispheres.

To further determine if overexpression of consecutively active Yki (Yki.S168A) causes a similar phenotype, we attempted to overexpress this protein using *TH-Gal4*. Similar to experiments performed using *pnr-Gal4,* this manipulation caused lethality, prohibiting us from performing HPLC on these animals. To overcome this issue, we performed a conditional overexpression experiment by integrating the *tub-Gal80[ts]* system. Surprisingly, overexpression of Yki.S168A in adulthood did not cause any increase in dopamine levels (**Fig S4B**). We also observed this when a stronger Yki GOF allele in which three key phosphorylation sites are mutated (Yki.S111A.S168A.S250A, a.k.a. Yki.S3A) [14] was overexpressed using the same paradigm (**Fig S4B**). We hypothesized that this may be because the Hippo pathway regulates dopamine levels only during development when pigmentation is actively taking place. To test this, we conditionally knocked down *wts* post-eclosion using *tub-Gal80[ts]* and also observed no difference in dopamine levels when this gene was manipulated only in adulthood (**Fig S4A**). In summary, we observe that inhibition of hippo signaling can modulate global dopamine levels during development but not post-development.

In flies, dopamine is synthesized as a precursor of melanin during development in epithelial cells and as a neuromodulator in dopaminergic neurons [50]. HPLC measurement of dopamine from whole fly heads theoretically reflects the total amount of this molecule made from both of these cell-types. Since we were interested in assessing whether Hippo signaling may regulate dopamine levels specifically in the brain, we next investigated whether key genes in the Hippo signaling pathway are expressed in dopaminergic neurons. We generated or obtained T2A-Gal4 gene trap lines that allow visualization of *wts* (*wts^T2A^*) and *yki* (*yki^T2A^*) expression *in vivo* [62,63] (**Fig 4C-4D**). We found that *wts* and *yki* are expressed in ∼50% or ∼75% of dopaminergic neurons we examined, respectively (**Fig 4C’-4D’**).

We next assessed whether knockdown of *wts* in dopaminergic neurons increase dopamine levels in the brain. To assess this, we manually dissected fly brains from *wts* KD flies and performed HPLC. Since *TH-Gal4* that is commonly used is known to miss a number of dopaminergic neurons, especially in the PAM (Protocerebral Anterior Medial) cluster [64], we utilized a recombined line in which *TH-Gal4* was combined with *GMRE5802-Gal4* [44,65], a transgenic construct in which Gal4 is expressed under the control of an enhancer element from DAT (*dopamine transporter*). In our previous study, we validated this line by knocking down *TH* and demonstrating that dopamine levels in dissected brains are significantly decreased [44]. Using this approach, we observed that brain dopamine levels were not changed using two independent *wts-RNAi* lines (**Fig 4B**). While this doesn’t rule out the possibility that defects in Hippo signaling may be causing a local alteration in dopamine levels, we did not obtain data that supports the role of this pathway in neuromodulation.

### snRNA-seq reveals multiple genes and pathways that contribute to Hippo signaling-mediated pigmentation phenotype

We next attempted to determine the genetic mechanism by which Hippo signaling regulates cuticle pigmentation. In a previous study, Oh et al, reported that *TH* and *Ddc* may be under the control of Yki based on Chromatin Immunoprecipitation sequencing (ChIP-seq) and bioinformatics studies [66]. In addition, True et al., reported that co-overexpression of TH and Ddc can darken the cuticle color in the fly wing [67]. To determine if this is also the case in the thorax, we performed overexpression and co-overexpression studies of TH and Ddc using *pnr-Gal4*. To our surprise, we found that neither TH overexpression nor TH/Ddc co-overexpression was sufficient to induce the dramatic cuticle pigmentation defects observed upon Hippo pathway inhibition (**Fig S5A-S5B**, **S5D**). Overexpression of *yellow* (*y*), which encodes an enzyme involved in melanogenesis, on its own as well as co-expression of *y* with *TH* also were also not sufficient to produce similar pigmentation defects (**Fig S5C, S5E**).

To determine how Hippo signaling regulates cuticle pigmentation, we decided to combine tissue specific RNAi experiments with transcriptomic analysis. Since the *pnr-Gal4* expressing epithelial cells are difficult to manually dissect out from other tissues (e.g. muscle, hemocyte, sensory neurons, trachea) in the developing fly thorax, we decided to perform snRNA-seq to determine the cell-autonomous effect of manipulating Hippo signaling specifically in epithelial cells (**Fig 5**). To distinguish *pnr-Gal4* expressing cells from non-*pnr* expressing cells, we used *UAS-CD8::GFP* alongside expressing a control RNAi or *wts-RNAi* (**Fig 5A**). We harvested pupa at 90 hours after puparium formation (hAPF) because key genes important in cuticle pigmentation have been documented to be highly expressed around this time point [68–70]. Through t-distributed stochastic neighbor embedding (t-SNE) analysis, we were able to identify many different expected cell types from the dissected tissue (**Fig 5B**), including a population of epidermal cells that were identified based on expression of *grainy head* (*grh*), a master regulator gene for epithelial cell identity and function [71,72]. We next focused on a subpopulation of these cells based on their expression of the UAS-transgenes, which allows us to identify the cells that express *pnr-Gal4* (**Fig S6A**). To further limit our analysis to epithelial cells that have initiated the cuticle pigmentation process, we further focused on a subset of *grh^+^ pnr-Gal4^+^*cells that also express *TH* for subsequent DEG (Differentially Expressed Gene) analysis. We found that there is noticeable separation between *control-RNAi* and *wts-RNAi* cells in a t-SNE plot (**Fig 5C**), indicating that they have some differences in their gene expression profiles. We identified 547 differentially expressed genes: 411 genes were significantly up-regulated (log_2_FC>0) and 136 genes were significantly down-regulated (log_2_FC<0) with an adjusted p-value of less than 0.05 and visualized these by volcano plot (**Fig 5D, Table S1A, S2A**). With a more stringent cut off of log_2_FC>0.5 (>∼41% increase) or log_2_FC<-0.5 (>∼29% decrease), we identify 364 differentially expressed genes; 310 upregulated and 54 downregulated. On this gene list, we performed a GO (gene ontology) analysis and found an enrichment for multiple biological terms that relate to Hippo signaling such as ‘positive regulation of hippo signaling’ (GO:0035332), ‘actin filament organization’ (GO:0007015), and ‘regulation of growth’ (GO:0040008) (**Fig S6B**, **Table S3**). Genes that are commonly used as readouts of Yki activity such as *expanded (ex)* and *Death-associated inhibitor of apoptosis 1* (*Diap1)* were identified to be significantly upregulated, as well as the long non-coding RNA, *CR43334*, which is the precursor of the micro-RNAi, *bantam* [73,74] (**Table S1A**). The upregulation of multiple known target genes of Hippo signaling confirms *wts* KD leads to increased Yki activity in the epithelial cells of interest.

**Fig 5.**
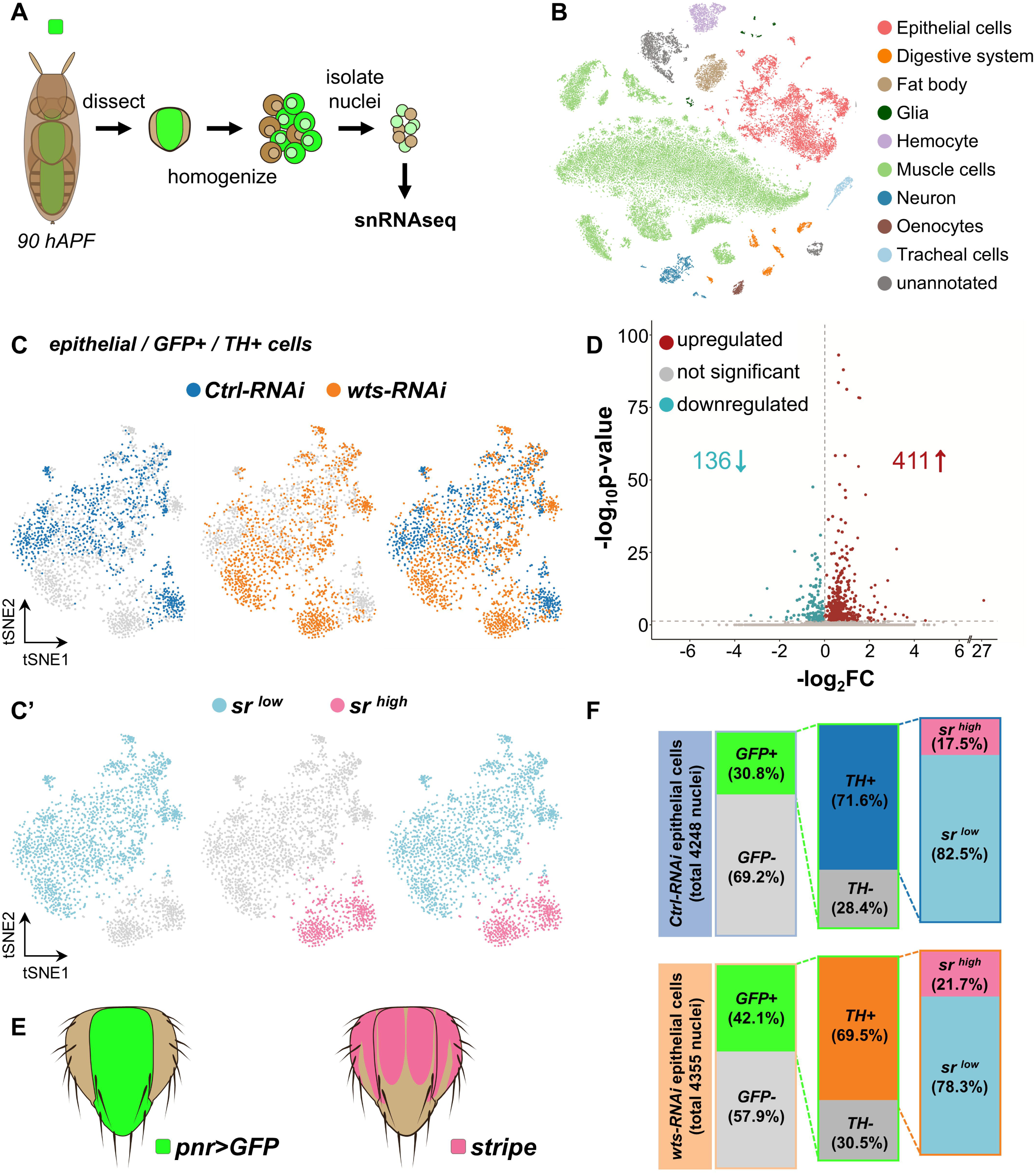
Cell populations and DEGs determined from *wts* RNAi pupal thorax by snRNA-sequencing. **(A)** Following the schematic, single nuclei were isolated, and transcripts were measured to produce a **(B)** t-=SNE plot to separate cells isolated from the notum by cell type. The cells of interest lay within the red indicated epithelial cluster (grey dashes) and were further visualized by **(C)** *pnr>CD8::GFP* expression and dopamine producing cells as indicated by the presence of *GFP* and *TH* transcripts. Control RNAi cells are shown in orange while *wts* RNAi KD cells are shown in blue. **(C’)** Epithelial cells were further subtyped into a *sr*^high^ expressing cell cluster in pink and *sr* ^low^ expressing cells in light blue, respectively. **(D)** Top genes with differential expression between control and *wts* KD cells from all *grh^+^/GFP^+^/TH^+^*cells are shown using a volcano plot. **(E)** Schematic of *pnr-Gal4* expression domain [147] in green and *stripe (sr)* expression domain [99] in pink based on previous studies. **(F)** Bar graphs representing cell population proportion based on *GFP^+^*, *TH^+^*, or *sr^high/low^*annotation from snRNA-seq datasets comparing Ctrl-RNAi to *wts*-RNAi.

Through this analysis, we also observed signatures that JNK (c-Jun N-terminal kinase) signaling, a pathway that is known to interact with the Hippo signaling pathway in multiple contexts [75–78], is activated upon *wts* KD. The second GO term enriched in our data set is ‘dorsal closure’ (GO: 0007391) (**Fig S6B**, **Table S3**), and this phenotype is commonly seem upon JNK signaling modulation [79,80]. The *pnr-Gal4* P-element insertion is known to mildly disrupt *pnr* expression and leads to a slight thorax closure phenotype at high temperatures, which seems to be exacerbated by *wts* KD (**Fig 1D, 1H, S7A-S7A’**). Genes in this ‘dorsal closure’ GO category as well as the greater DEG list as a whole include many upregulated JNK related genes such as; *canoe* (*cno*) [81,82], *cryptocephal* (*crc*) [83], *Diap1* [84], *flapwing* (*flw*) [85], *happy hour* (*hppy*) [86], *misshapen* (*msn*) [87,88], *Protein tyrosine phosphatase 61F* (*Ptp61F*) [89], *PDGF- and VEGF-receptor related* (*Pvr*) [80], *Ras-like protein A* (*Rala*) [90], *raw* [91,92] and ultimately the JNK signaling activity reporter gene, *puckered* (*puc*) [93–95] (**Table S1A**). Interestingly, a JNK signaling activator, *Ask1 (Apoptotic signal-regulating kinase 1)*, was previously been found to modulate cuticle pigmentation in the thorax [96,97]. To determine if activation of JNK signaling contributes to the phenotypes seen with Hippo signaling inhibition, we knocked down *basket* (*bsk,* the JNK encoding gene) as well as *hemipterous* [*hep,* a JNK kinase (JNKK) encoding gene] to inhibit JNK signaling activity (**Fig S7B-S7C**). Although we did not observe any major changes in the clefting phenotype mediated by *wts* KD (**Fig S7A’-S7C’**) and the notum size is not modulated (**Fig S7D**), inhibition of JNK signaling significantly suppressed the pigmentation defects (**Fig S7A-C’, S7E-S7F**). This indicates that modulation of JNK signaling is one mechanism by which Hippo signaling regulates cuticle pigmentation.

Additionally, we found that altered expression of a transcription factor that plays a role in epidermal cell identity may also contribute to the altered pigmentation phenotype upon Hippo signaling manipulation. Epithelial cells in the notum can be classified into cells that do or do not express *stripe* (*sr*) [98,99] (**Fig 5C’**, **5E**). Cells that expresses Sr, a zinc finger transcription factor, are called tendon cells, and LOF mutations in this gene causes increased melanization in a portion of the notum [99]. Because we noticed that the increase of pigmentation in our *wts* KD nota seems to follow the *sr* expression pattern (**Fig 1C-H, 5E, S7E**), we decided to further classify the *grh^+^ pnr-GAL4^+^ TH^+^* cells based on *sr* expression (**Table S1B-C, S2B-C**). In our snRNA-seq dataset, *sr*^high^ and *sr*^low^ expression cells clearly segregate on a t-SNE plot (**Fig 5C’**), indicating that tendon and non-tendon epithelial cells have different gene expression programs. While we isolated a similar number of total cells (Ctrl RNAi; 23,052 cells v.s. *wts* RNAi; 23,138 cells) and epithelial cells (Ctrl RNAi; 4,248 v.s. *wts* RNAi; 4,355) for each genotype, there was a greater ratio of *pnr* driven GFP vs non-GFP cells in *wts* KD (42.1%) compared to control (30.8%) (**Fig 5F, S8A**). Within these groups, *TH* expression followed a similar ratio in control (71.6%) vs wts KD (69.5%), but upon *wts* KD, there is a slight increase in the proportion of *sr^high^* cells (*wts* KD 21.7% vs Ctrl 17.5%). In addition, when focusing on these *Grh^+^ pnr-Gal4^+^ TH^+^ sr^high^* epithelial cells, we noticed that *sr* transcripts themselves were significantly reduced in our *wts* KD group compared to control (**Table S1B**). Together, loss or reduction of *sr* may also contribute to the increased pigmentation as well as the specific pattern observed upon Hippo signaling manipulation.

Since tendon cells and non-tendon cells may have different gene regulatory networks that responds slightly different to manipulation of Hippo signaling, we a further performed DEG analysis on *sr^high^* and *sr^low^* cells, separately (**Table S1B-C, Table S2B-C**). We identified 77 DEGs that are specific to *sr^high^* cells and 11 DEGs that are specific to *sr^low^* cells (**Fig S8B**). While the directionality of DEGs were consistent in most DEGs identified, we found 3 instances where DEGs (*CG4962*, *how*, *shot*,) show opposite pattern of regulation between these two cell-types (**Fig S8B**). This suggests that while Hippo signaling modulates similar sets of genes between tendon and non-tendon cells, there are few targets genes that are regulated differently between these two types of epithelial cells.

In order to identify additional genes that work downstream of *wts* and contribute to the cuticle pigmentation defect observed upon KD of this gene, we next attempted to identify genetic modifiers of the dark cuticle phenotype based on epistatic RNAi analyses. To prioritize the genes to test experimentally, we decided to focus on DEGs which have been previously linked to pigmentation phenotypes in the literature (**Fig 6A**). We first crossed referenced all significant DEGs we found from the snRNA-seq experiment (**Fig S8B, Table S1A-C**) against genes which have been annotated with the phenotypic term ‘Abnormal Body Color (FBcv:0000356)’ in FlyBase (flybase.org; total 526 genes curated as of June 2025) and identified 53 pigmentation candidate genes that may function work downstream of *wts*. We next verified the information in FlyBase by manually reading the referenced manuscripts to confirm that manipulation of these genes have been reported to alter body color [44,56,99–109]. For 46 of these genes, increase or decrease in pigmentation was reliably reported in the notum, abdomen and/or wing. Additionally, we independently found three DEGs in which pigmentation phenotypes were reported in the literature but not annotated with the ‘Abnormal Body Color’ terminology in Flybase (ie. *Duox* [110], *InR* [111] *and sd* [107,109]). We next asked how many of these total 49 genes (**Fig S8C**, **Table S1**) have a consistent phenotype with the expression change in our data set. Concordance was determined by identifying DEGs with increased expression that show a pale phenotype upon LOF or darkened phenotype upon GOF. DEGs with decreased expression that were reported to show a darkened phenotype upon LOF were also considered to be concordant. Of the 49 confirmed pigmentation genes, 24 genes have concordant phenotypes while 25 have non-concordant phenotypes. Based on availability of reagents to manipulate these genes, we tested the effect of RNAi-mediated KD of 14 concordant and 11 non-concordant genes to determine whether they suppress or enhance the *wts* KD pigmentation phenotype (**Fig 6, S9**). Expression of all of these genes were increased upon *wts* KD so we hypothesized that KD of concordant genes that contribute to the increased pigmentation will behave as genetic suppressers whereas KD of some non-concordant genes may act as genetic enhancers.

**Fig 6.**
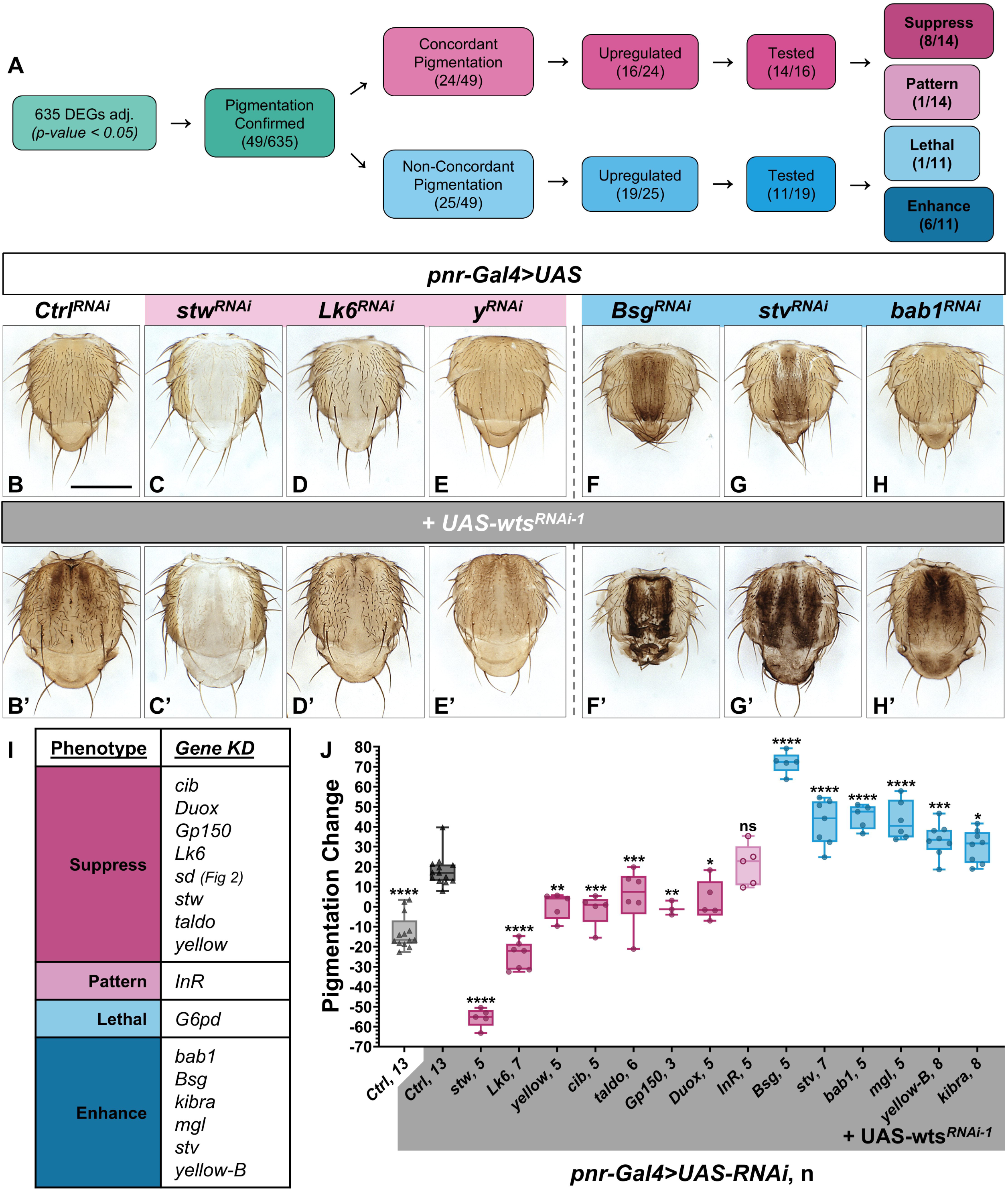
Cumulative effects of pigmentation regulating DEGs are responsible for increased pigmentation phenotype upon *wts* knockdown. **(A)** To narrow down the list of genes for further functional screening, DEGs from snRNA-seq analysis were prioritized based on previous literature and available RNAi reagents. **(B)** Compared to control RNAi, (C) *stw-*RNAi KD has a strong pale phenotype on it’s own **(C’)** and is able to suppress the pigmentation phenotype in a (B’) *wts*-RNAi KD background. **(D/D’)** KD of *Lk6* on its own shows a pale phenotype as well and can also suppress the wts KD pigmentation. **(E/E’)** KD of *y* on its own has no pigmentation phenotype in the notum but is also able to suppress *wts* KD pigmentation. RNAi KD of (F/F’) *Bsg,* (G/G’) *stv,* and (H/H’) *bab1* all have growth and darkening phenotypes are their own and **(F’-H’)** enhance the darkened cuticle upon co-KD with *wts*. Growth temperature is 29°C and scale bar = 0.5mm. **(I)** Table of determined category of modulation for each interacting DEG. **(J)** Plotted grey value difference between *pnr* domain of scutum and internal tissue control to determine pigmentation change. Statistical significance was determined by One-Way ANOVA pair-wise analysis compared to *pnr>wts-RNAi; Ctrl-RNAi* (n=3-13, 29°C). Precise method of pigmentation analysis can be found in **Fig S7E** while quantification data for growth phenotypes can be found in **Fig S9K**.

We were able to suppress the increased cuticle pigmentation phenotype of *wts* KD by targeting 8 of the 14 concordant genes [e.g. *straw* (*stw*), *Lk6* kinase (*Lk6*), *yellow* (*y*), *ciboulot* (*cib*), *transaldolase* (*taldo*), *Gp150*, *Dual oxidase* (*Duox*) and *sd*] (**Fig 2J-2K’, 6B-6E’,6I-J**, **S9A-S9E’**). In addition, consistent with previous reports, knockdown of *InR* alone led to reduced pigmentation in the nota [111], and co-KD of *InR* and *wts* exhibited a change in pigmentation pattern, even though average pigmentation intensity was unchanged compared to *wts* KD alone (**Fig 6I-J, S9F-S9F’**, **S9J-K**). Of the 9 genes that modified the pigmentation phenotype induced by *wts* KD, only 2 genes, *InR* and *sd*, significantly suppressed tissue size defects (**Fig 2J-2K’, 6J, S2C, S9F-S9F’**, **S9K**). While we observed a reduction in nota size when *cib*, *taldo,* and *Gp150* were knocked down alone, co-KD of these genes as well as with *stw*, *Lk6*, *y* or *Duox*, had no significant impact on the *wts* KD induced tissue overgrowth phenotype (**Fig S9K**).

We were also able to observe enhancement of the pigmentation phenotype seen upon *wts* KD by co-KD of 6 out of the 11 non-concordant genes tested [e.g. *Basigin* (*Bsg*), *starvin* (*stv*), *bric à brac 1* (*bab1*), *Megalin* (*mgl*), *yellow-B*, and *kibra*] (**Fig 6F-6I**, **S9G-S9I’**). KD of *Glucose-6-phosphate dehydrogenase* (*G6pd)* with our without *wts* RNAi led to lethality (**Fig 6I**). On their own, KD of *Bsg*, *stv*, *bab1* and *yellow-B* reduces the size of the nota, but only co-KD of *Bsg* was able to suppress the *wts* KD overgrowth phenotype (**Fig S9K**). Importantly, *kibra*, a known target gene and positive regulator of Hippo signaling that contributes to a negative feedback loop of this pathway [112,113], is upregulated in our dataset. KD of *kibra* alone led to increased nota size as well as slightly increased pigmentation [56] and significantly enhanced both the growth and pigmentation phenotype upon co-KD with *wts* (**Fig 6I-6J**, **S9I-S9I’**, **S9K**). These observations are consistent with a model in which a negative feedback loop exists to fine tune Hippo signaling outputs, and some of the DEGs we observed may be upregulated as a compensatory mechanism to prohibit further hyperpigmentation and/or growth upon *wts* KD.

## Discussion

In this study, we report a novel role of the canonical Hippo signaling pathway in cuticle pigmentation in *Drosophila*. To date, most studies regarding Hippo signaling pathway have focused on its role in growth and development because the primary phenotype one observes when this pathway is manipulated in mosaic animals is overgrowth or undergrowth of tissues [7,36–39]. One reason that this pigmentation phenotype has been under-appreciated is likely because most of these studies have used scanning electron microscopy (SEM) which lack information on color. Although we can appreciate that *wts* and *mats* mutant clones generated in a few previous studies exhibit pigmentation defects that are strikingly similar to what we document in this study based on the images provided in three manuscripts [36,37,39], these authors did not further investigate this phenotype. In this study, we convincingly show that LOF of all four kinases of the Hippo signaling pathway (*hpo*, *sav*, *mats*, *wts*) give a similar increased cuticle pigmentation phenotype, which is also seen upon overexpression of the activated Yki protein (**Fig 1**, **2F-2F’**, **S2B**). We further show that the increased pigmentation phenotype caused by *wts* KD is dependent on Yki as well as Sd (**Fig 2G-2K’**), which are the main transcriptional factors that mediate the canonical Hippo signaling. Interestingly, Sd was also identified as a modulator of abdominal pigmentation in two studies that investigated gene regulatory mechanisms of abdominal pigmentation in *Drosophila* [107,109]. Therefore, our data agrees with scattered evidence in the literature that canonical Hippo signaling mediated by Yki and Sd regulates cuticle pigmentation. Since cuticle pigmentation plays critical roles in fitness such as adaptation [114], resistance to environmental stressors [115–117] and ultimately natural selection [53,118], further studies regarding how modulation of this pathway in this essential biological process may provide important insights into evolution.

During development, cuticle pigmentation is regulated by multiple enzymes including TH and Ddc (**Fig S1A**), which also play critical roles in dopamine synthesis in the brain [50,119,120]. By comparing the level of dopamine in the head and brain through HPLC analysis, we found that *wts* does not seem to have a major role in regulating global dopamine levels in the brain despite being expressed in many dopaminergic neurons (**Fig 4A-4D**). We have recently shown that while some genes that affect pigmentation affect dopamine levels in both fly heads and brains, the majority of the genes that have strong pigmentation defects do not affect brain dopamine levels when knocked down in dopaminergic cells [44]. More specifically, in an HPLC-based screen of 35 genes that affect cuticle pigmentation, we found 11 genes that affect head dopamine levels. However, only two (*clueless* and *mask*) of these eleven genes were found to have significant effects on total brain dopamine levels. This may suggest that mechanisms that regulate dopamine levels in epithelial cells are different from mechanisms that govern dopamine levels in the brain. Indeed, different splicing isoforms of *TH* and *Ddc* are expressed in the epithelial cells and in dopaminergic neurons [121,122], suggesting that different gene regulatory networks of dopamine synthesis, at least at the level of mRNA splicing, have developed over evolution. This is further demonstrated by our finding that overexpression of the cuticle isoform of TH, which strongly increases dopamine levels in the fly head, is not capable of doing so when expressed in dopaminergic neurons in the brain (**Fig 4B**). Since cuticle color and behavior are both under selective pressure in the wild, there may have been some benefits in establishing different mechanisms to control these two phenomena that involve dopamine separately [123].

By performing snRNA-seq on pupal nota, we identified >600 genes that were significantly differentially regulated upon *wts* KD. Manual inspection of these DEGs (e.g. *ex*, *diap1, CR34443/bantam,* and *kibra*) (**Table S1**) as well as GO analysis (**Fig S6B, Table S3**) consistently showed that our experimental manipulation is effectively inhibiting the Hippo signaling pathway. A majority of the DEGs were upregulated, which is also consistent with our understanding that canonical Hippo signaling primarily inhibits transcription by sequestering Yki activity. Interestingly, while previous studies suggested that *TH* and *Ddc* may be under the transcriptional control of Yki [66], we did not find these genes to be significantly altered in the contexts we analyzed.

Through epistatic analysis, we determined that JNK signaling contributes to Hippo signaling mediated pigmentation. Expression of *puc,* a standard readout of JNK activity [93–95], is significantly increased upon *wts* KD, and suppression of JNK activity through *bsk* or *hep* co-KD suppressed the *wts* induced pigmentation defect without affecting the overgrowth phenotype (**Figure S7**). Prior to this work, one study linked an upstream component of JNK signaling, Ask1, to cuticle melanization in the fly nota [84,96]. In contrast to our study, however, this manuscript concluded that while Ask1 can act as a JNKK kinase (JNKKK), this gene primarily acts via the Mkk4(MAP kinase kinase 4)-p38 pathway, rather than through the classical JNK axis that involves Hep and Bsk, to induce melanization. Hence, our finding identifies cuticle pigmentation as a novel context in which Hippo and JNK signaling intersect biologically.

Another potential functional link between Hippo and JNK signaling can be postulated based on one DEG we functionally investigated as a modulator of *wts* KD pigmentation, *yellow-b*. Although *yellow-b* has not been linked to cuticle pigmentation phenotypes before this study, it is a member of the *yellow* gene family [124,125]. KD of *yellow-B* shows no pigmentation change in the nota but can significantly enhance the *wts* KD pigmentation phenotype (**Fig 6J, S9H-S9H’**). This was included as a non-concordant gene in our screen list as it is an enhancer of the *wts* KD phenotype and is upregulated, rather than being downregulated, upon *wts* KD.

Interestingly, *yellow-b* expression has been reported to be altered when JNK signaling is activated in a variety of contexts [126,127], possibly through transcriptional regulation mediated by AP-1 (Activator Protein-1, a dimer formed by c-Jun and c-Fos) [128]. Hence, our study provides a list of genes that may help dissect the complex relationships between Hippo signaling and JNK signaling, both of which are relevant to a variety of pathogenic conditions such as congenital disorders and cancer.

Based on snRNA-seq analysis, we also found that *TH* expressing epithelial cells in the developing fly nota transcriptionally segregate into two major subpopulations based on expression levels of Sr in t-SNE clustering. We were able to identify 363 DEGs in our *wts* KD population compared to control in the *sr^high^* group while there were 362 DEGs in the *sr^low^*population, 206 of which were shared between the two cell types (**Fig S8B, Table S1B**). Interestingly, we found that the *sr* transcript levels was significantly decreased in the *sr^high^* cell population upon *wts* KD while ration of *sr^high^* to *sr^low^*epithelial cells didn’t change dramatically. As Sr itself is a negative regulator of melanization [99] and it’s expression pattern is reflective of the *wts* KD pigmentation increase pattern (**Fig 5E**, **6B’**), disruption of tendon cell identity may contribute to increase in pigmentation, which warrants further investigations. By mining previously published transcriptomic datasets, we found two previous studies that also show that *sr* transcripts are reduced in the wing discs of a *wts* mutant [66] or upon overexpression of *yki.S168A* [129]. These observations indicate that Hippo signaling may directly or indirectly regulate Sr levels in different developmental contexts.

To identify additional genes that are regulated by Hippo signaling and contribute to cuticle pigmentation, we further narrowed down our combined list of DEGs for experimental validation and performed genetic epistasis analysis for 25 genes (**Fig 6A**). Through this effort, we identified eight genes that contribute to pigmentation increase, one that modifies the pigmentation pattern, and six that potentially serve to dampen the severity of pigmentation increase (**Fig 6I**). In the following section, we elaborate on how some of these factors may function downstream of Hippo signaling in the context of pigmentation and growth regulation.

Two concordant DEGs, *stw* and *y*, encode enzymes involved in melanization (**Fig S1A**) and behave as suppressors of the *wts* KD pigmentation phenotype (**Fig 6C’-6D’**). Interestingly, the cis-regulatory region of *y* has been previously found to be bound by Sd through a yeast-one hybrid approach [107], indicating that *y* may be a direct target of Hippo signaling. Additionally, one of the non-concordant DEGs found in our list that enhanced the pigmentation phenotype upon *wts* co-KD, *mgl*, has previously been found to regulate endocytic clearance of Y protein during wing cuticle formation [106]. When *mgl* levels are reduced, Y protein levels increase and lead to increased melanization in the wing. If this function is conserved in the dorsal throax, increase of *mgl* transcripts may serve as a negative feedback mechanism to prevent hyperpigmentation. Another gene found in our DEG list that has been studied in the context of wing pigmentation is *Duox*. Previously, it was reported that *Duox* KD in the posterior wing region, lead to decreased pigmentation and fragility in the wings [110]. This study proposed that Duox plays a role in reactive oxygen species (ROS) production and increased ROS in this context destabilizes the wing by decreasing catechol content and tyrosine cross-linking. This can theoretically affect both sclerotization and melanization needed for proper wing development. The gene *stw* mentioned above encodes Laccasse 2, an enzyme that is also known to be required for both sclerotization and melanization by direct synthesis of quinones needed for these processes [106] (**Fig S1**). Therefore, the upregulation of both *Duox* and *stw* upon *wts* KD may indicate that Hippo signaling may also affect sclerotization in addition to melanization which would require further investigation.

Identification of *kibra* as a non-concordant genetic enhancer of both pigmentation and growth phenotypes of *wts* KD highlights the importance of a negative regulatory mechanism that is built into the Hippo signaling pathway [112,113]. While KD of *kibra* alone shows a slight increase in pigmentation and significant increase in nota size (**Fig S9I-S9I’, S9K**), co-KD of *kibra* and *wts* enhanced both the growth and pigmentation than either KD alone (**Fig 6B’, 6J**). Increased *kibra* expression in the scRNA-seq dataset is evidence that inhibition of Hippo signaling through *wts* KD is capable of activating a transcriptional feedback loop and additional DEGs identified through our transcriptomic analysis may help identify additional factors that function as regulatory safeguards. Bab1, identified as a non-concordant enhancer from our study, may serve in this role. This transcrion factor has been proposed to function to as a central player in mediating thermal plasticity of pigmentation in *Drosophila* [130] and has also been studied as a mediator of sexual dimorphic pigmentation traits, especially in the fly abdomen [103]. Future studies focused on this and other transcription factors identified as DEGs upon Hippo signaling modulation may help refine the complex negative as well as positive feedback loops that finetune this important signaling pathway.

Importantly, five genes we identified as suppressors or enhancers (*cib, Gp150, Lk6, taldo*, *stv*) of the hyperpigmentation phenotype observed upon *wts* KD have unknown molecular roles in cuticle pigmentation as they were only identified in large scale screens as pigmentation alleles [44,56]. Identification of these genes as Hippo signaling pigmentation modifiers using independent reagents used in this study further confirms their role as *bona fide* cuticle pigmentation regulators. Interestingly, all 14 of the DEG pigmentation modifiers found in this study have been identified to be bound by Yki and/or Sd or reported to be regulated by Hippo signaling in different contexts [66,107,129,131]. While direct molecular validation is needed, some of these genes may turn out to be underappreciated direct targets of Hippo signaling and warrant further mechanistic investigations.

It is worth noting that were not able to classify *InR* as a suppressor or enhancer of *wts* RNAi-mediated pigmentation phenotype based on the pigmentation quantification method that we developed for this study. We find that KD of *InR* is able to partially suppress the overgrowth phenotype and modulates the pigmentation pattern in the fly nota without affecting the average pigmentation intensity upon co-KD with *wts* (**Fig S9F-S9F’, S9J-S9K**). This finding is consistent with a previous study that reported that nutritional status can affects both growth and pigmentation through Insulin/IGF (Insulin-like growth factor) and TOR (target of rapamycin) pathways in *Drosophila* [111], and some genes may have context specific roles (e.g. may have different downstream effect in *sr^high^*and *sr^low^* cells) in altering pigmentation patterns. Since our functional screen to follow up on the DEGs were limited to 24 genes, further genetic studies to characterize the role of >600 genes that are significantly altered upon Hippo signaling inhibition in the developing pupal nota will likely lead to discovery of understudied genes in the context of cuticle pigmentation, tissue growth, or both.

Finally, *Drosophila melanogaster* has become a major model organism to study the functional consequences of rare genetic variants identified in individuals with rare genetic disorders [132] or cancer [133,134]. Disruptions of the Hippo signaling pathway have widespread effects on human health across development, physiology and oncogenesis of multiple organ systems [17–35, 40–43]. For example, recessive variants in *STK4* (one of two human orthologs of *hpo*) cause ‘Immunodeficiency 110 with Lymphoproliferation (MIM# 614868)’ [135,136]. In addition, dominant *YAP* (one of two human orthologs of *yorkie*) variants is known to cause ‘Coloboma, ocular, with or without hearing impairment, cleft lip/palate, and/or impaired intellectual development (MIM#120433)’ [137]. Additionally, rare variants in *LATS1* (one of two human ortholog of *wts*) have been identified in a case of familial cerebral cavernous malformations[138], as well as in urinary bladder and colon cancer [139]. While bioinformatic tools can be used to predict whether the variant may alter gene or protein function of rare variants found in patients, experimental studies are often needed to determine the precise functional consequences of VUS (variants of uncertain significance) [140]. In a previous study, homozygous lethality of *wts* mutant was shown to be fully rescued by human LATS1 [141]. Consistently with this finding, we show that cuticle pigmentation and growth phenotypes of multiple genes can be rescued by their human orthologs (MOB1A/B, LATS1/2 and YAP1, Fig 3 and S3). Because both LOF and GOF of Hippo signaling can cause readily identifiable scorable phenotypes in the notum, this platform could be used as a convenient assay system to study the functional consequences of VUS in canonical Hippo signaling genes in rare disease patients or in cancer. In summary, the underappreciated role of canonical Hippo signaling that we discovered here could have direct translational application in genomic medicine.

## Materials and Methods

### Fly maintenance

Fly (*Drosophila melanogaster*) lines were maintained at room temperature on a molasses-based food source in vials or bottles. Crosses were set with the same food plus sprinkled yeast to promote egg laying, typically with 3-4 males and 5-8 virgin females per tube, reared mainly at 29°C on a 12:12hr Light/Dark (LD) program and transferred every three to four days (unless stated otherwise). Additional temperatures used for crossing are stated otherwise and include 25°C and 18° until 3^rd^ instar larvae stage or after eclosion, then transferred to 29°C to make use of the temperature sensitive expression of the Gal4/UAS system as well as the temperature sensitive Gal80[ts]. For proper aging, eclosed flies were gathered 0-3 days after eclosion (DAE) into separate tubes and aged 1-5 DAE for notum imaging and 3-7 DAE for gene expression analysis and HPLC measurement.

### Fly stocks

Publicly available fly lines were obtained from Bloomington *Drosophila* Stock Center (BDSC, https://bdsc.indiana.edu/), Vienna Drosophila Research Center (VDRC, https://www.viennabiocenter.org/vbcf/vienna-drosophila-resource-center/), or the Japanese National Institute of Genetics (https://shigen.nig.ac.jp/fly/nigfly/). Lines that are not publicly available were gifted from other labs or produced for this manuscript. A list of all fly lines used can be found in **Table S4**. Information on DEG RNAi lines tested with co-KD of *wts* that produced no obvious modulation of pigmentation or growth phenotypes can found in **Table S5**.

### UAS-human cDNA overexpression construct and transgenic *Drosophila* generation

Transgenic UAS-Human cDNA fly lines were generated following previous protocols [142]. In brief; UAS-Human cDNA constructs were generated through Gateway cloning to produce an LR reaction between donor and destination plasmids [143]. Donor human cDNA plasmids were provided from The Kenneth Scott Collection from the Department of Molecular and Human Genetics at Baylor College of Medicine. The following transgenic plasmid constructs were generated based (donor plasmid, GenBank, clone); pGW.UAS-MOB1A.attB (pDONR221, NM_018221.3, IOH26027), pGW-MOB1B.attB (pDONR221, NM_018221.3, IOH3008), pGW.UAS-LATS1.attB (pDONR221, NM_004690.2, IOH45203), pGW.UAS-LATS2.attB (pENTR233.1, NM_014572.2, KSID65), pGW.UAS-YAP1.attB (pDONR221, NM_001130145.1, IOH26027). Transgenic flies were generated by UAS-Human cDNA integration into the VK37 docking site via a □C31 integrase reaction [144].

### Generation of *wts-T2A-Gal4* line

The *wts^T2A^* allele used was generated by ΦC31-mediated recombination-mediated cassette exchange of a MiMIC (Minos mediated integration cassette) insertion line [145]. Conversion of the original MiMIC element (MiMIC: 05605) was performed by genetic crossing of UAS-2xEGFP, hs-Cre,vas-dΦC31, Trojan T2A-Gal4 triplet flies to the MiMIC strain and following a previously established crossing scheme [146].

### Notum dissection, imaging, and quantification

For RNAi KD or overexpression (OE) cuticle phenotype analysis, *y w; pnr-Gal4/TM3,Sb*[147] or *pnr-Gal4,UAS-TH-RNAi/TM3,Sb* virgins were crossed to *UAS-RNAi/OE* males and kept at 29°C until 1-5 DAE unless temperature is stated otherwise. For somatic CRISPER knockout (sKO) experiments, virgins of the genotypes *UAS-Cas9; pnr-Gal4/TM3,Sb* or *UAS-Cas9; pnr-Gal4,UAS-TH-RNAi/TM6b* were crossed to males ubiquitously expressing guide RNAs at 29°C until 1-5 DAE. For temperature-controlled RNAi KD or OE expression analysis, *tubGal80[ts]; pnr-Gal4/TM6b,Tb* or *tubGal80[ts]; pnr-Gal4,UAS-TH-RNAi /TM3,Sb* virgins were crossed to *UAS-RNAi/OE* males at 18°C until being switched to 29°C at the 3rd larvae instar stage until 1-5 DAE. Rescue or suppression of *wts-RNAi* LOF was performed by crossing *wts-RNAi/(FM7,Kr::GFP); pnr-Gal4/TM3,Sb* virgins to *UAS-RNAi/OE* males at the temperature stated until 1-5 DAE. For direct pigmentation analysis, flies were aged matched to account for aging differences in pigmentation. Flies were stored in 70% EtOH at room temperature until dissection and imaging.

Using an adapted protocol from [44,148], nota were dissected and imaged. The adapted protocol is as follows; thoraces were dissected in 70% EtOH by removing in order the head, wings, abdomen and legs, ensuring there was a hole on the ventral side leading to the abdominal cavity for 10% KOH to reach the soft tissues. The thoraces were put into 100μl of 10% KOH and heated on a block for 10 minutes at 85°C. The KOH was then removed, and the thoraces were washed 3 times in 500μL of 70% EtOH. Each notum was trimmed out in 70% EtOH, removing the extra tissue from the ventral side along the natural divot between dorsal and ventral side of the thorax. For mounting, the nota were first placed into mounting media (50% glycerol and 50% 190 Proof EtOH) to displace any bubbles on the nota. Then the nota were transferred to a drop of this same media on a slide with 2 layers of white tape (VWR Tape, #89097-986) on both sides to raise the coverslip to allow for the height of the thorax. To capture z-stack brightfield images we used a Leica MZ16 stereo microscope with an OPTRONICS MicroFIRE camera or Leica Z16 APO stereo microscope with an DMC4500 Digital Microscope camera to capture z-stack brightfield images. These images were combined using extended depth of field in Image-Pro Plus 7.0 with In-Focus (v1.6) and Leica Application Suite X (v3.7.6.25997), respectively. For each genotype, 3-13 different nota were imaged per genotype, with the best representative image used in final figures (brightness adjusted to normalize backgrounds). A document that contains all images acquired for quantification can be downloaded as **Extended Supplemental Data**.

#### Notum Size Quantification

ImageJ [149] was used to quantify the size of the *pnr-Gal4* expressing area as well as the area of the entire notum to get a read-out of the *pnr* proportion of notum. For each image, the ImageJ freehand selection tool was used to outline the *pnr* domain only to capture the size of this area, and then another measurement was taken outlining the entire notum. The *pnr* area was then calculated as a portion of the entire notum by using the equation pnr area/notum area. This final value was plotted using GraphPad Prism 10 to visualize each genotype compared against control/genotype indicated in the figure. Using GraphPad, a one-way ANOVA was run to determine statistical significance against indicated controls because data sets were normally distributed and had equal standard deviations. When comparing only two groups, an Unpaired t-test was performed to determine significance. P-values represented with asterisks are as follows; ****<0.0001, ***<0.001, **<0.01, and *<0.05.

#### Notum Pigmentation Plot Profiles and Quantification

ImageJ [149] was also used to create plot profiles and quantify pigmentation of the nota by way of calculating the grey values of each notum and subsection of notum. For pattern comparison, a Grey Value Plot Profile of a section of the notum encompassing both side control tissues and the *pnr-Gal4* expressing area, was created by drawing a rectangle across the middle of the notum just above the macrochaetae bristles (indicated in green in **Fig S9J**). For wild-type sized controls, this rectangle was 0.525mm by 0.3mm while the nota with an overgrowth phenotype was increased to 0.7mm by 0.35mm. The nota with *InR-RNAi* growth rescue wts-RNAi KD was measured by 0.525mm by 0.35mm to capture the proportionally same area. The resulting individual Grey Values were then subtracted from the Grey Value of the background of the image (measured using a 0.2mm by 0.2mm square). The values were then plotted using Microsoft Excel with a line connecting the individual points to visualize the pigmentation pattern changes across the notum. For easier visualization on the full plot profile across the notum, the blue block corresponds to control tissue, and pink corresponds to the *pnr* expressing domain excluding the midline that is shown with white (**Fig S9J-J’**).

To quantify the pigmentation changes for statistical analysis, the freehand selection tool of ImageJ was again used to draw along the pnr domain expressing border of the scutum only to obtain the mean grey value of the area (pink outline in **Fig S7E**) The grey values for the control tissue, not expressing *pnr-Gal4*, on the sides of the nota was captured with a rectangle measuring 0.3mm by 0.1mm for control wild type sized flies and 0.35mm by 0.1mm co-KD *wts* nota (in blue **Fig S7E**). For the final grey value difference for plotting in GraphPad Prism 10, the *pnr-Gal4* expressing area value (pink) was subtracted from the average of the control tissue grey value (blue, **Fig S7E**). Using GraphPad, a one-way Anova was run to determine statistical significance against indicated controls because data sets were normally distributed and had equal standard deviations. P-values represented with asterisks are as follows; ****<0.0001, ***<0.001, **<0.01, and *<0.05.

### Brain expression analysis of *wts* and *yki*

*UAS-mCherry::nls (mCh::nls)* virgins were crossed to either *wts^T2A^/TM3,Sb* or *yki^T2A^/SM6a* males at 25°C and aged 3-7 DAE for dissection similar to Marcogliese et al., 2022 [142]. Fly brains were dissected in ice cold 1X PBS and fixed in 4% PFA (diluted with 0.5% PBST) rocking at 4°C for ∼18hrs in a 24 well plate. Brains were then washed 2X quickly and then 3X rocking for 15min each in 0.5% PBST (these washing steps were repeated after both the primary and secondary incubations). The PBST was replaced with primary antibody consisting of 1:500 anti-Tyrosine Hydroxylase-Rabbit (Pel-Freez Biologicals, P40101) diluted in 0.5% PBST and rocked for 2 days at 4°C. The washing PBST was then replaced by the secondary antibody consisting of 1:200 anti-Rabbit-Alexa647 (Thermo Fisher Sci., A-27040) diluted in 0.5% PBST, for two hours at room temperature. After washing in PBST, same as between the fixing and primary antibody incubation, the brains were mounted in Vectashield mounting media (H-1900) and imaged on a Zeiss LSM 710 confocal microscope using z-stacks throughout the entire brain. The Zeiss ZEN software was used to create the Z-Projection images seen here. For quantification of co-expression of mCh::nls and TH, we manually counted each dopaminergic cluster per hemisphere (PAL, PPL1, PPL2ab, PPM2/3) and represented this as a percentage of number of mCh::nls positive and TH positive cells devided by the total number of TH positive cells per dopaminergic cluster. The total fractions of dopaminergic neurons expressing each genes were calculated by adding all clusters together for a final value. The graphs to visualize these results were made using GraphPad Prism 10.

### High Performance Liquid Chromatography (HPLC) analysis of dopamine

For head expression analysis, *TH-Gal4* (*ple-Gal4*, #8848) virgins were crossed to *UAS-RNAi/OE* males and reared at 29°C until 3-7 DAE. For temperature-controlled head expression, *tubGal80[ts];TH-Gal4* virgins were crossed to *UAS-RNAi/OE* males at 18°C until eclosion and then raised at 29°C until 3-7 DAE. For brain measurements *TH-Gal4,GMR58E02-Gal4*[44] were crossed with *UAS-RNAi/OE* males at 29°C until 3-7 DAE. The following protocol was adapted from previous studies [44,150,151]

#### Sample Preparation

To measure head dopamine concentrations, female heads were collected with a razor blade under CO2 anesthetic during the time period ZT02-ZT07. 60µL of 50 mM citrate acetate (pH=4.5) was added to each sample (n=5 heads per sample, 4-23 samples per genotype) and samples that weren’t analyzed same day were stored at −20°C. To measure brain concentration, flies were reared at 29°C until 3-7 DAE then put on ice for dissection. After removing the brains from flies in ice cold 1X PBS, brains were added to 60 µL of 50 mM citrate acetate (pH=4.5) on ice and stored at −20°C if not analyzed the same day (n= 5 female brains and 5 male brains, 4-7 samples per genotype).

For HPLC analysis, samples were ground with a pestle using a Cordless Pestle Motor and Fisherbrand Disposable Pellet Pestle for 1.5mL tube for 3 bouts of ∼20s seconds each. To get rid of debris, samples were spun for 10 minutes at 13,000 RPM and supernatant removed into a 300µL Polypropylene Sample Vial with 8mm Snap Caps. For heads only, 10 µL from each sample was used for the Bradford Protein Analysis Assay. The remaining sample was used for HPLC analysis.

#### Bradford Protein Analysis Assay

For protein measurement, the Bio-Rad Bradford Assay for colorimetric scoring of total protein was used (Bio-Rad Protein Assay Kit I #5000001). The dye reagent was diluted 1:4 in MilliQ H2O before being filtered through a 0.22 µm SFCA Nalgene filter. Using a protein standard from the kit, standards were diluted to protein concentrations of 500 µg/mL, 250 µg/mL, 100 µg/mL, and 50 µg/mL. In a 96-well plate, 10 µL of each protein standard or fly sample was placed into single wells along with 200 µL of the diluted 1:4 dye reagent. The samples incubated at room temperature for ∼30 minutes and then absorbance was measured using a BMG Labtech FLUOstar OPTIMA microplate reader. Final protein measurements for each sample were calculated based on standards ran on the same plate.

#### HPLC Information

The HPLC used is a Antec Scientific product with a LC110S pump, SYSTEC OEM MINI Vacuum Degasser, AS110 autosampler, a SenCell flow cell with salt bridge reference electrode in the Decade Lite. The column used was an Acquity UPLC BEH C18 Column (130Å, 1.7 µm, 1 mm X 100 mm with Acquity In-Line 0.2 µm Filter). The mobile phase was a 6% Acetonitrile mobile phase optimized for our samples and degassed for 10 minutes using the Bransonic Ultrasonic Bath (74.4 mg NA2EDTA·2H20, 13.72 mL 85% w/v phosphoric acid, 42.04 g citric acid, 1.2 g OSA, 120 mL acetonitrile, H20 up to 2L, pH=6.0 using 50% NaOH solution). Data was collected and processed using the DataApex Clarity chromatography software.

The standards for HPLC were made fresh the day of HPLC analysis, by diluting master stocks of 100 mM dopamine (Sigma-Aldrich, Cat#H8502) and 10 mM serotonin (Sigma-Aldrich, Cat#H7752), diluted in MilliQ H2O and stored at 4°C. Standards were diluted to concentration of 5, 10, 50, and 100 nM for dopamine and serotonin. Final sample concentrations of dopamine and serotonin were calculated based on standards run in the same day/batch. Standards were loaded into 300 µL Polypropylene Sample Vials with 8mm Snap Caps for HPLC analysis.

#### Statistical Analysis for HPLC

Dopamine measurements (pg/sample) were first normalized to protein concentration (µg/mL) of the same sample. This value was then normalized to the mean value of same day controls’ measurements. Statistical analysis was performed using Graphpad Prism 10. All data was subjected to a ROUT outlier test, where all outliers were removed. A one-way Anova was run for data sets with Gaussian distribution and equal standard deviations. Data sets without Gaussian distribution were analyzed via Kruskal-Wallis pair-wise comparison while data sets with Gaussian distribution but unequal standard deviation were analyzed via Welch’s ANOVA pair-wise comparison. Additionally, samples were normalized to the mean measurement of the controls run on the same day as well the amount of protein in the sample to account for differences in homogenization. P-values represented with asterisks are as follows; ****<0.0001, ***<0.001, **<0.01, and *<0.05.

### snRNA-seq of dissected pupal nota

Single nuclei RNA-sequencing experiments were performed by manually dissecting the pupal thorax, isolating single nuclei, preparing the library and conducing short-read sequencing based on the following steps.

#### Sample preparation for snRNA-seq

In bottles, using 10 males of the UAS-RNAi lines (control; *UAS-lacZ-RNAi*[152] and experimental; *UAS-wts-RNAi* [12702R-1]) and ∼30 virgin females (*pnr-Gal4,UAS-CD8::GFP/TM3, Sb*) crosses were set at 25°C. White pupae were moved to separate tubes at 25°C and after aging for 90 hours, dissected. 90 hAPF), pupae were placed on double sided tape on a slide, dorsal side up. First the operculum was removed and then slits made down the left and right sides of the pupal casing, the pupal casing was pulled back to expose the anterior half of the pupa. The thorax was removed from the head and abdomen and then the ventral half of the thorax was removed including the legs and wings. The dorsal half of the thorax was placed immediately into a 1.5ml RNAase free Eppendorf tube on dry ice until flash-frozen using liquid nitrogen. 20 thoraces of each genotype were used for each sample and stored at −80°C until further preparation for snRNA-seq.

#### Library preparation and sequencing for snRNA-seq

Single-nucleus suspensions were prepared following the protocol described previously with the adaptation of 20 strokes with the loose Dounce pestle and 20 strokes with the tight Dounce pestle [153]. Next, we used the BD AriaIII FACS sorter to collect nuclei. Nuclei were stained by Hoechst-33342 on ice (1:1000; >5min). Hoechst+ nuclei were collected during sorting. Individual nuclei were collected into one 1.5ml RNase-free Eppendorf tube with 200µl 1x PBS with 0.5% BSA as the receiving buffer (RNase inhibitor added). For each 10x Genomics run, 100k nuclei were collected. Nuclei were spun down for 10 min at 950g at 4°C and then resuspended using 80µl or desired amount of 1x PBS with 0.5% BSA (RNase inhibitor added). 2µl of nucleus suspension was used for counting the nuclei with hemocytometers to calculate the concentration. We loaded 40K nuclei to the 10x controller to target >20k nuclei for each channel.

Next, we performed snRNA-seq using the 10x Genomics platform with the Chromium Next GEM Single Cell 3’ HT (high-throughput) Reagent Kits v3.1 (Dual Index) with the following settings. All PCR reactions were performed using the BioRad C1000 Touch Thermal cycler with a 96-deep Well Reaction Module. The recommended cycle numbers from the 10x protocol were used for cDNA amplification and sample index PCR. As per the 10x protocol, 1:10 dilutions of amplified cDNA and final libraries were evaluated on a bioanalyzer. The final library was sent to Novogene Corporation Inc. for Illumina NovaSeq PE150 S4 lane sequencing with the dual index configuration Read 1 28 cycles, Index 1 (i7) 10 cycles, Index 2 (i5) 10 cycles, and Read 2 90 cycles. A PhiX control library was spiked in at 0.2 to 1% concentration. The sequencing depth is about 26-30K reads per nucleus.

#### snRNA-seq data processing

Raw snRNA-seq data, in the form of FASTQ files, underwent alignment to the *Drosophila melanogaster* reference genome (FlyBase release 6.31 with GFP sequence included) using the Cell Ranger software (v7.2.0). Subsequent steps involved the removal of ambient RNA contamination via CellBender [154] and the identification and exclusion of potential doublet cells using Scrublet [155]. Quality control criteria necessitated the elimination of cells exhibiting fewer than 200 genes or 500 UMIs. Genes detected in fewer than three nuclei were removed from our analysis. Furthermore, cells with gene or UMI counts exceeding five median absolute deviations from the median were also excluded from the analysis. Additionally, cells harboring over 5% of mitochondrial transcripts were eliminated. The majority of the snRNA-seq data analysis was conducted using the Scanpy package (v1.9.6, [156]).

#### Cell type annotation from snRNA-seq data

Our approach to annotating cell types closely mirrored the methodology previously established for AFCA annotations [157]. We integrated the thorax data with existing AD-FCA dataset [158], facilitating their co-clustering. To mitigate batch effects and align dataset variations, the Harmony algorithm [159] was applied to the co-clustered data. Subsequent to adjustment, AD-FCA-derived cell type labels were assigned to thorax cells using a Logistic Regression classifier, with AD-FCA serving as the training set and thorax dataset as the test set. These initial automated annotations were subsequently subjected to manual validation and correction to enhance reliability. We sub clustered epithelial cells to annotate GFP+ epithelial cells. And further we defined TH (*ple*)+ cells within GFP+ epithelial cells for further analyses. Additionally, based on the Leiden clustering of total epithelial cells (resolution = 0.5), two distinct clusters exhibiting enriched expression of *stripe* (*sr*) were designated as *sr*^high^ cells, while the remaining clusters were characterized as *sr*^low^ cells for further analysis. Raw FASTQ files, expression matrix, and processed h5ad files, including cell type annotations, are available from NCBI/GEO (accession number GEO: GSE310450).

#### DEG and gene ontology analysis

To identify genes with altered expression levels in *wts-RNAi* flies compared to *LacZ-RNAi* controls, we performed DEG analysis using the Wilcoxon Rank Sum test. Genes were considered differentially expressed if they met a false discovery rate (FDR) threshold of <0.05.

Differential expression analysis yielded genotype-specific DEGs, encompassing both upregulated and downregulated genes. These DEGs were subjected to GO analysis using the GOATOOLS software (v1.2.3)[160]. For this purpose, the gene association dataset (FB2025_07) was retrieved from FlyBase, with a specific focus on Biological Process (BP) GO terms for our investigations.

## Acknowledgements

We thank the Bloomington *Drosophila* Stock Center (USA), the National Institutes Genetics Fly Stock Center (Japan), and the Vienna Drosophila RNAi Center (Austria) for providing useful fly stocks and reagents for this project. We also thank FlyBase for providing genetic and genomic data, infrastructure, and tools essential to this study. We would like to thank Drs. Herman Dierick, Sheng Zhang, Hugo Bellen, Oguz Kanca, Michael Wangler, Huda Zoghbi, and Daryl Scott for useful suggestions, discussions and advice on this project. We also thank Dr. Hirokazu Hashimoto, Ms. Mei-Chu Huang, Ms. Hongling Pan, and Ms. Danqing Bei for their efforts in establishing some of the transgenic fly lines used in this work. This work was supported by startup funds to SY from the Jan and Dan Duncan Neurological Research Institute at Texas Children’s Hospital (NRI Fellowship) and the Department of Molecular and Human Genetics at Baylor College of Medicine (SEED Funds). SBG was supported by The Cullen Foundation. Confocal microscopy at BCM was supported in part by the Intellectual and Developmental Disabilities Research Center (IDDRC, U54HD083092) from the Eunice Kennedy Shriver National Institute of Child Health & Human Development. Generation of UAS-human cDNAs transgenic lines was supported by the NIH (R24OD022005 to SY). This project was also supported by the Cytometry and Cell Sorting Core at Baylor College of Medicine with funding from the CPRIT Core Facility Support Award (CPRIT-RP240432), the NIH (P30CA125123 and S10OD036336) with the assistance by Mr. Joel M. Sederstrom. The funders had no role in study design, data collection and analysis, decision to publish, or preparation of the manuscript.

## Author Contributions

S.Y., S.B.G., S.L.D., and J.W.M. conceived the experiments. S.Y. and S.B.G wrote the manuscript. S.B.G. and H.C. created transgenic lines for this paper. S.B.G. conducted all nota, head, and brain preparation for HPLC and brightfield or confocal imaging. S.B.G. and J.W.M. conducted image analysis. S.B.G. and S.L.D conducted the HPLC analysis. S.B.G, Y.J.P. and Y.Q. conducted snRNA-seq experiments. S.B.G., Y.J.P., B.S. and H.L. conducted data analysis for snRNA-seq. All authors participated in the critical analysis of the manuscript.

## Supplemental Figure Titles and Legends

**Fig S1. Inhibiting the dopamine synthesis pathway results in pale cuticle. (A)** Core dopamine metabolism and melanin synthesis pathway in *Drosophila melanogaster* (adapted from Deal et. al [44]). **(B)** Dissected control notum compared to **(C)** pale cuticle phenotype seen in a *pnr>TH-RNAi* knockdown notum. Performed at 29°C and scale bar = 0.5mm.

**Fig S2. Quantification of nota size for Hippo signaling transcriptional effectors and epistasis. (A)** *yki* OE enhances the severe phenotype seen upon Hippo signaling kinase KD (*hpo* or *wts*). Statistical significance was determined by one-way ANOVA pair-wise compared to *yki* OE (n=3-5, 25°C). **(B)** Overgrowth observed upon *tub-Gal80^[ts]^; pnr>yki.S168A* expressed from the 3^rd^ larval instar (LI) stage (n=5-6, 18>3^rd^ LI>29°C) Statistical significance was determined by an unpaired t-test. **(C)** Undergrowth of *pnr>sd*-*RNAi* KD persists even with addition of *wts*-*RNAi* KD. Statistical significance was determined by one-way ANOVA pair-wise analysis compared to *Ctrl*s. (n=4-10, 25°C).

**Fig S3. Quantification of overgrowth rescue mediated by human ortholog overexpression. (A)** Expression of either *MOB1A* and *MOB1B* rescues overgrowth phenotype observed with *pnr>mats-RNAi KD*. Statistical significance was determined by one-way ANOVA pair-wise compared to *Ctrl*s (n=4-8, 29°C). **(B)** Expression of either *LATS1* and *LATS2* rescues overgrowth phenotype observed with *pnr>wts-RNAi KD*. Statistical significance was determined by one-way ANOVA pair-wise compared to *Ctrl*s (n=3-10, RT).

**Fig S4. Adult stage manipulation of Hippo signaling genes shows no significant change in head dopamine levels. (A)** Using *tub-Gal80[ts]; ple-Gal4>UAS*, head DA levels were assessed by HPLC for *wts* KD in the adult stage and showed no difference. (n=6-11, 5 heads/n, 18>AE>29°C, 3-7 DAE). Statistical significance was determined by Kruskal-Wallis pair-wise compared to control. **(B)** Head DA levels do not change when *yki.SA* is overexpressed in the adult stage (n=19-24, 5 heads/n, 18>AE>29°C, 3-7 DAE). Statistical significance was determined by Brown-Forsythe and Welch’s ANOVA pair-wise compared to control **(C)** KD of *yki* by RNAi in the adult stage shows no changes in head DA levels (n=4-10, 5 heads/n, 18>AE>29°C, 3-7 DAE). Statistical significance was determined by one-way ANOVA compared to control.

**Fig S5. Minimal pigmentation differences observed when overexpressing multiple melanin synthesis genes. (A/B)** *pnr>control* OE compared to **(B)** *TH* or **(C)** *y* OE shows minimal pigmentation increase. **(D)** Co-expression of *TH* and *Ddc* or **(E)** *TH* and *y* with *pnr>Gal4* does not show small pigmentation increase from *TH* or *y* OE on its own. All performed at 29°C. Scale bar = 0.5mm.

**Fig S6. snRNA-seq of *pnr>GFP* epithelial cells upon Hippo signaling modulation. (A)** All *GFP* positive cells in every cell cluster type with quantified representation of *GFP*, *grh*, and *Hml* expression in each cell cluster type. **(B)** GO terms of DEGs with adjusted p-value < 0.05.

**Fig S7. Hippo signaling regulates pigmentation through active JNK signaling.** Compared to **(A)** Ctrl-RNAi, **(B)** *bsk*-RNAi KD, **(C)** *hep*-RNAi and **(A’)** *wts*-RNAi KD produce slight cleft nota phenotypes. Inhibiting JNK signaling through **(B’)** *bsk* KD or **(C’)** *hep* KD does not affect **(D)** nota size. **(E)** To measure pigmentation change, the mean grey scale value of the *pnr* domain of the scutum (pink) was subtracted from the mean grey value of the internal control tissue (blue). **(F)** Inhibition of JNK signaling via *bsk* or *hep* KD suppress the pigmentation increase seen with *wts* KD. Growth temperature is 29° C. Scale bar = 0.5mm. Statistical significance was determined by One-Way ANOVA pair-wise compared to *pnr>wts-RNAi; Ctrl-RNAi* (n=5-13, 29°C).

**Fig S8. Subclassification of cells based on *stripe* (*sr*) expression allows for DEG analysis of different epithelial cell subtypes. (A)** Table of total cell counts for each gene expression condition with proportion of total population in brackets. **(B)** Venn diagram showing the overlap between different clustering of cells based on All grh^+^/GFP^+^/TH^+^ vs sr^high^/grh^+^/GFP^+^/TH^+^ vs sr^low^/grh^+^/GFP^+^/TH^+^. Three DEGs, found to show differential regulation between the cell types are documented in a small table. **(C)** Venn diagram when only comparing validated pigmentation related gene expression overlaps. Number of DEGs increased indicated with an up arrow while decrease is shown with a down arrow.

**Fig S9. While certain screen hits change overall pigmentation with or without affecting growth, *InR* changes pigmentation patterning and suppresses growth.** Additional DEGs screened for pigmentation and growth modulation include; **(A)** Compared to control **(B)** *cib-*RNAi KD has a slight pale phenotype on it’s own **(B’)** and is able to suppress the pigmentation phenotype in a (A’) *wts*-RNAi KD background. **(C/C’)** KD of *taldo* on its own shows a slight pale phenotype and also suppresses wts KD pigmentation **(D/D’)** KD of *Doux* on its own has no pigmentation phenotype in the notum but is also able to suppress *wts* KD pigmentation. RNAi KD of **(E/E’)** *Gp150* has a slight pale phenotype and suppresses *wts* KD pigmentation. **(F/F’)** *InR* KD is the only concordant gene manipulation that suppresses *wts* KD growth and shows now overall change in mean pigmentation. **(G)** *mgl* has no growth or pigmentation phenotype on its own, **(G’)** but suppresses pigmentation of *wts* KD. **(H)** *yellow-B* KD has a growth but no pigmentation phenotype while **(H’)** co-KD with *wts* shows an enhanced pigmentation. **(I)** KD of *kibra* shows slight darkening phenotype and increased notum size, while enhancing both the growth and pigmentation phenotype with wts KD. Scale bar = 0.5mm. **(J)** Schematic of notum areas analyzed to determine grey values corresponding to pigmentation intensity. The green blocked area was used to visualize pigmentation intensity of **(J’)** individual plot across the nota based on its inverse grey value. Corresponding *pnr* area (pink) and control area (blue) to further subdivide nota areas for visualization. Growth temperature is 29°C. Statistical significance was determined by One-Way ANOVA pair-wise analysis. If indicated with grey asterisks or “ns” below box-plot, values compared to *pnr>Ctrl-RNAi and* if indicated with black asterisks or “ns*”* above box plot, values compared to *pnr>wts-RNAi; Ctrl-RNAi* (n=3-13, 29°C). Details of the quantification method can be found in the **Materials & Methods** section and **Fig 1I**.

## Supplemental Table Titles and Legends

**Table S1A. List of significantly differentially expressed genes (DEGs) upon *wts* KD in TH expressing epithelial cells.** Specifics on DEGs (log2FC >0 or <0 and adjusted p-value<0.05). Associated phenotypes for pigmentation related genes found through FlyBase “abnormal body color” allele curation, FlyBase “cuticle pigmentation” GO terms, or independent literature search.

**Table S1B: List of significantly differentially expressed genes (DEGs) upon *wts* KD in *TH*^+^ *sr*^high^ expressing epithelial cells.** Specifics on DEGs (log2FC >0 or <0 and adjusted p-value<0.05). Associated phenotypes for pigmentation related genes found through FlyBase “abnormal body color” allele curation, FlyBase “cuticle pigmentation” GO terms, or independent literature search.

**Table S1C. List of significantly differentially expressed genes (DEGs) upon *wts* KD in *TH*^+^ *sr*^low^ expressing epithelial cells.** Specifics on DEGs (log2FC >0 or <0 and adjusted p-value<0.05). Associated phenotypes for pigmentation related genes found through FlyBase “abnormal body color” allele curation, FlyBase “cuticle pigmentation” GO terms, or independent literature search.

**Table S2A. Genes detected by sn-RNA sequencing in TH expressing epithelial cells, *wts* KD vs *Ctrl*.** All genes with a log_2_FC>0 or <0 but not all statistically significant changes.

**Table S2B. Genes detected by sn-RNA sequencing in *TH* expressing epithelial cells, *sr*^high^, *wts* KD vs *Ctrl*.** All genes with a log_2_FC>0 or <0 but not all statistically significant changes.

**Table S2C. Genes detected by sn-RNA sequencing in *TH* expressing epithelial cells, *sr*^low^, *wts* KD vs *Ctrl*.** All genes with a log_2_FC>0 or <0 but not all statistically significant changes.

**Table S3. Top GO Terms for DEGs.** DEGs with a log2FC<0.5 or <0.5 and associated GO term enrichmenents for biological processes, cellular components and molecular functions. Based on FlyBase GO terms (FB2025_07).

**Table S4: List of transgenic *Drosophila* lines used and additional information.**

**Table S5: *Drosophila* lines that produced no obvious change of *wts* KD phenotypes.**

