## Supplementary material for "Hippo signaling regulates cuticle pigmentation and dopamine metabolism in *Drosophila*": Table S4

**Table S4: List of transgenic *Drosophila* lines used and additional information.**

| LINE | GENOTYPE | SOURCE | STOCK NUMBER |
|---|---|---|---|
| pnr-Gal4 | *yw;P{GawB}*$pnr^{MD237}$*/TM3,Sb* | BDSC<br>*rebalanced with TM3,Sb | 3039 |
| pnr-Gal4, UAS-TH-RNAi | *yw;P{GawB}*$pnr^{MD237}$*,P{TRiP.JF01813}attP2/TM3,Sb* | This study<br>(recombined BDSC 3039 and BDSC 25796) | |
| TH-Gal4 | $w^{*}$*; P{ple-GAL4.F}3* | BDSC | 8848 |
| R58-Gal4 | $w^{1118}$*; P{GMR58E02-GAL4}attP2* | BDSC | 41347 |
| TH-Gal4, R58-Gal4 | *yw; P{ple-GAL4.F}3, P{GMR58E02-GAL4}attP2* | Deal et. al [44] | |
| tub-Gal80$^{ts}$ | *w*; P{tubP-GAL80[ts]}10; TM2/TM6B,*$Tb^{1}$ | BDSC | 7108 |
| tub-Gal80$^{ts}$; pnr-Gal4 | *w*; P{tubP-GAL80[ts]}10/CyO; P{GawB}*$pnr^{MD237}$*/TM6B,*$Tb^{1}$ | This study<br>(double balanced BDSC 7108 and BDSC 3039) | |
| tub-Gal80$^{ts}$; pnr-Gal4, UAS-TH-RNAi | *w*; P{tubP-GAL80[ts]}10/CyO; P{GawB}*$pnr^{MD237}$*,P{TRiP.JF01813} attP2/TM3,Sb* | This study<br>(double balanced BDSC 7108 and recombined BDSC 3039 and BDSC 25796) | |
| UAS-Cas9; pnr-Gal4 | *w*; P{UAS-Cas9.P2}attP40/ CyO;P{GawB}*$pnr^{MD237}$*/TM6B,*$Tb^{1}$ | BDSC | 67077 |
| UAS-Cas9; pnr-Gal4, UAS-TH-RNAi | $y^{1}$*w*; P{UAS-Cas9.P2}attP40/CyO; P{GawB}*$pnr^{MD237}$*,P{TRiP.JF01813} attP2/TM3,Sb* | This study | |
| wts[TG4] | $y^{1}$*w*; Mi{Trojan-GAL4.0} wts[MI05605-TG4.0]/TM3,Sb* | This study | |
| yki[TG4] | $y^{1}$*w*;TI{GFP[3xP3.cLa]=CRIMIC.TG4.1}yki[CR00812-TG4.1]/SM6a* | BDSC | 79628 |
| pnr-Gal4, UAS-CD8::GFP | $y^{1}$*w*; pnr-Gal4, P{w[+mC]=UAS-mCD8::GFP.L}LL6/TM3,Sb* | This study<br>(recombined BDSC 3039 and BDSC 5130) | |
| UAS-Ctrl-RNAi | *w P{*$w^{+}$*, UAS>lacZ.SL.IR}12a* | Kennerdell and Carthew [152] | |
| UAS-TH-RNAi | $y^{1}v^{1}$ *; P{TRiP.JF01813}attP2* | BDSC | 25796 |
| UAS-hpo-RNAi$^{1}$ | $y^{1}v^{1}$ *; P{TRiP.HMS00006}attP2* | BDSC | 33614 |
| UAS-hpo-RNAi$^{2}$ | $y^{1}v^{1}$ *; P{TRiP.JF02740}attP2* | BDSC | 27661 |
| UAS-hpo-RNAi$^{3}$ | $w^{1118}$; P{GD1570}v7823 | VDRC | v7823 |
| UAS-mats-RNAi$^{1}$ | $y^{1}sc^{*}v^{1}sev^{21}$ *; P{TRiP.HMS00475}attP2* | BDSC | 34959 |
| UAS-mats-RNAi$^{2}$ | $y^{1}sc^{*}v^{1}sev^{21}$ *; P{TRiP.JF03246}attP2* | BDSC | 29567 |
| UAS-mats-RNAi$^{3}$ | $w^{1118}$*; P{GD8136}v17716* | VDRC | v17716 |
| UAS-wts-RNAi$^{1}$ | *P{NIG.12072R}2* | NIG-Japan | 12702R-2 |
| UAS-wts-RNAi$^{2}$ | *P{NIG.12072R}1* | NIG-Japan | 12702R-1 |
| UAS-wts-RNAi$^{3}$ | $w^{1118}$*; P{GD1563}v9928* | VDRC | v9928 |
| UAS-yki-RNAi$^{1}$ | *P{NIG.4005R}3* | NIG-Japan | 4005R-3 |
| UAS-yki-RNAi$^{2}$ | $w^{1118}$*; P{GD11187}v40497/TM3* | VDRC | v40497 |
| UAS-sd-RNAi$^{1}$ | $y^{1}v^{1}$*; P{TRiP.JF02514}attP2* | BDSC | 29352 |
| UAS-sd-RNAi$^{2}$ | $y^{1}sc^{*}v^{1}sev^{21}$ *; P{TRiP.GL00410}attP2* | BDSC | 35481 |
| UAS-bsk-RNAi | $y^{1}$ $v^{1}$*; P{TRiP.JF01275}attP2* | BDSC | 31323 |
| UAS-hep-RNAi | $y^{1}sc^{*}v^{1}sev^{21}$ *; P{ TRiP.GL00089}attP2* | BDSC | 35210 |
| UAS-stw-RNAi | $w^{1118}$*; P{GD12744}v22959* | VDRC | v22959 |
| UAS-Lk6-RNAi | $w^{1118}$*; P{GD9353}v32885* | VDRC | v32885 |
| UAS-yellow-RNAi | $y^{1}sc^{*}v^{1}sev^{21}$ *; P{TRiP.HMC05546}attP40* | BDSC | 64527 |
| UAS-Bsg-RNAi | $w^{1118}$*; P{GD15718}v43306* | VDRC | v43306 |
| UAS-stv-RNAi | $w^{1118}$*; P{GD10796}v34408* | VDRC | v34408 |

| | | | |
|---|---|---|---|
| UAS-bab1-RNAi | $y^1 sc^* v^1 sev^{21}$; *P{TRiP.HMC04716}attP40* | BDSC | 57410 |
| UAS-cib-RNAi | $y^1 v^1$; *P{TRiP.JF02837}attP2* | BDSC | 28003 |
| UAS-taldo-RNAi | *P{NIG.2827R}2* | NIG-Japan | 2827R-2 |
| UAS-Duox-RNAi | $y^1 sc^* v^1 sev^{21}$; *P{TRiP.HMS00692}attP2* | BDSC | 32903 |
| UAS-Gp150-RNAi | *P{NIG.5820R}2* | NIG-Japan | 5802R-2 |
| UAS-InR-RNAi | $y^1$ $v^1$; *P{TRiP.JF01183}attP2* | BDSC | 31594 |
| UAS-mgl-RNAi | $y^1$ $v^1$; *P{TRiP.JF02545}attP2* | BDSC | 27242 |
| UAS-yellow-B-RNAi | *P{NIG.17914R}1* | NIG-Japan | 17914R-1 |
| UAS-kibra-RNAi | $y^1$ $v^1$; *P{TRiP.JF03098}attP2* | BDSC | 28683 |
| EGFP-gRNA | $y^1 sc^* v^1 sev^{21}$; *P{TKO.GS00382}attP40* | BDSC | 79393 |
| hpo-gRNA | $y^1 sc^* v^1 sev^{21}$; *P{TKO.GS04500}attP40* | BDSC | 85878 |
| sav-gRNA | $y^1 sc^* v^1 sev^{21}$; P{TKO.GS05550}attP40/CyO | BDSC | 84080 |
| UAS-LacZ | *w*; P{UAS-lacZ.Exel}2* | BDSC | 8529 |
| UAS-TH | *w*; P{UAS-ple.T}331f2* | BDSC | 37539 |
| UAS-yki.S168A$^1$ | $y^1 w^*$; *P{UAS-yki.S168A.GFP.HA}10-7-Y* | BDSC | 28816 |
| UAS-yki.S168A$^2$ | $w^*$; *P{UAS-yki.S168A.V5}attP2* | BDSC | 28818 |
| UAS-yki.S3 | *w*; P{UAS-yki.S111A.S168A.S250A.V5}attP2* | BDSC | 28817 |
| UAS-MOB1A | *yw; pGW.UAS-MOB1A* | This study | |
| UAS-MOB1B | *yw; pGW.MOB1B* | This study | |
| UAS-LATS1 | *yw; pGW.UAS-LATS1* | This study | |
| UAS-LATS2 | *yw; pGW.UAS-LATS2* | This study | |
| UAS-yki | *w*; P{UAS-yki.V5.O}attP2* | BDSC | 28819 |
| UAS-YAP1 | $y^1 w^*$; *pGW.UAS-YAP1* | This study | |
| UAS-yellow | $y^1 w^{1118}$; *P{UAS-y.C}MC1* | BDSC | 3043 |
| UAS-Ddc | $w^*$; *P{UAS-Ddc.T}16f2/TM3,Sb*$^1$ | BDSC | 37540 |
| UAS-TH, UAS-Ddc | $y^1 w^*$; *P{UAS-ple.T}331f2,P{UAS-Ddc.T}16f2/TM3, Sb* | This study (recombined BDSC 37539 and BDSC 37540) | |

*BDSC=Bloomington Drosophila Stock Center, VDRC=Vienna Drosophila Resource Center, NIG= National Institute of Genetics
