## Supplementary material for "Hippo signaling regulates cuticle pigmentation and dopamine metabolism in *Drosophila*": Table S5

**Table S5: *Drosophila* lines that produced no obvious change of *wts* KD phenotypes.**

| DEG PHENOTYPE | LINE | GENOTYPE | SOURCE | STOCK NUMBER |
|---|---|---|---|---|
| Concordant | UAS-CG46639 | $y^{1}sc*v^{1}sev^{21}$ ; *P{ TRiP.HMS01636}attP2* | BDSC | 37494 |
| Concordant | UAS-Hrb98DE-RNAi$^{1}$ | $w^{1118}$; *P{GD14939}v29523* | VDRC | v29523 |
| Concordant | UAS-Hrb98DE-RNAi$^{2}$ | *P{NIG.9983R}4* | NIG-Japan | 9983R-4 |
| Concordant | UAS-Mp-RNAi$^{1}$ | $y^{1}v^{1}$; *P{ TRiP.JF02929}attP2* | BDSC | 28299 |
| Concordant | UAS-Mp-RNAi$^{2}$ | $y^{1}v^{1}$; *P{TRiP.HMJ21668}attP40* | BDSC | 52981 |
| Concordant | UAS-Mp-RNAi$^{3}$ | *P{NIG.8647R}2* | NIG-Japan | 8647R-2 |
| Concordant | UAS-Mp-RNAi$^{4}$ | *P{NIG.8647R}3* | NIG-Japan | 8647R-3 |
| Concordant | UAS-sima$^{1}$ | $y^{1}v^{1}$; *P{TRiP.JF02105}attP2* | BDSC | 26207 |
| Concordant | UAS-sima$^{2}$ | $y^{1}sc*v^{1}sev^{21}$ ; *P{TRiP.HMS00832}attP2* | BDSC | 33894 |
| Concordant | UAS-sima$^{3}$ | *P{KK102226}VIE-260B* | VDRC | v106187 |
| Concordant | UAS-Synd$^{1}$ | $y^{1}v^{1}$; *P{ TRiP.JF02607}attP2/TM3, Sb[1]* | BDSC | 27297 |
| Concordant | UAS-Synd$^{2}$ | $w^{1118}$; *P{GD9087}v40018* | VDRC | v40018 |
| Concordant | UAS-Synd$^{3}$ | *P{KK109082}VIE-260B* | VDRC | v104580 |
| Non-Concordant | UAS-cdi-RNAi$^{1}$ | $y^{1}v^{1}$; *P{ TRiP.JF03006}attP2* | BDSC | 28369 |
| Non-Concordant | UAS-cdi-RNAi$^{2}$ | $y^{1}v^{1}$; *P{TRiP.HMJ02227}attP40* | BDSC | 42568 |
| Non-Concordant | UAS-cdi-RNAi$^{3}$ | $y^{1}sc*v^{1}sev^{21}$ ; *P{TRiP.HMS04453}attP40/CyO* | BDSC | 57010 |
| Non-Concordant | UAS-cdi-RNAi$^{4}$ | $y^{1}sc*v^{1}sev^{21}$ ; *P{TRiP.GL01860}attP2/TM3, Sb[1]* | BDSC | 57032 |
| Non-Concordant | UAS-CG42671-RNAi$^{1}$ | $y^{1}sc*v^{1}sev^{21}$ ; *P{TRiP.GL01098}attP2* | BDSC | 36844 |
| Non-Concordant | UAS-CG42671-RNAi$^{2}$ | *P{KK104428}VIE-260B* | BDSC | v100467 |
| Non-Concordant | UAS-CG42674-RNAi$^{1}$ | $y^{1}v^{1}$; *P{TRiP.JF01659}attP2/TM3, Sb[1]* | BDSC | 31166 |
| Non-Concordant | UAS-CG42674-RNAi$^{2}$ | $y^{1}sc*v^{1}sev^{21}$ ; *P{TRiP.HMS00228}attP2* | BDSC | 34943 |
| Non-Concordant | UAS-Vrp1-RNAi$^{1}$ | $y^{1}sc*v^{1}sev^{21}$ ; *P{TRiP.HMS01593}attP2* | BDSC | 36704 |
| Non-Concordant | UAS-Vrp1-RNAi$^{2}$ | $y^{1}sc*v^{1}sev^{21}$ ; *P{ TRiP.GL00640}attP40* | BDSC | 38201 |

*BDSC=Bloomington Drosophila Stock Center, VDRC=Vienna Drosophila Resource Center, NIG= National Institute of Genetics
